# Responsive fluorophore aggregation provides spectral contrast for fluorescence lifetime imaging

**DOI:** 10.1101/2020.01.28.923672

**Authors:** Kelton A. Schleyer, Benjamin D. Datko, Brandon Burnside, Chao Cui, Xiaowei Ma, John K. Grey, Lina Cui

## Abstract

Fluorophores experience altered emission lifetimes when incorporated into and liberated from macromolecules or molecular aggregates; this trend suggests the potential for a fluorescent, responsive probe capable of undergoing self-assembly and aggregation and consequently altering the lifetime of its fluorescent moiety to provide contrast between the active and inactive probes. We developed a cyanobenzothioazole-fluorescein conjugate (**1**), and spectroscopically examined the lifetime changes caused by its reduction-induced aggregation *in vitro*. A decrease in lifetime was observed for compound **1** in a buffered system activated using the biological reducing agent glutathione, suggesting a possible approach for designing responsive self-aggregating lifetime imaging probes.

## Introduction

The lifetime of a fluorescence event is an intrinsic property of the fluorescent species in question. The same fluorophore can possess unique lifetime values if its population is in two distinct environments, resulting in contrast created by the same reporter^1, 2^. Lifetime can be altered chemically, such as by coordination with metal ions^3, 4^ or through acid-base equilibrium with environmental pH^5, 6^; it is also exquisitely sensitive to small changes in environmental factors like temperature^7^ and solvent viscosity^8, 9^. The lifetime of a fluorophore is influenced by its capacity to undergo non-radiative emission^10^, which is altered by immobilizing factors such as tethering to macromolecules^11^, entrapment in aggregated proteins^12^, or intercalation into DNA^13^. In these examples, the lifetime of the free fluorophore is altered when exposed to a macromolecular aggregate.

Fluorescein isothiocyanate (FITC) has served as a reporter in several lifetime studies involving aggregation or macromolecules. When bound to the active site of the anti-fluorescein antibody Mab 4-4-20, FITC was observed to have a major lifetime component of 0.37 ns, compared to the 3.86 ns major component observed for the fluorophore free in solution^14^. A poly-L-lysine oligomer bearing multiple FITC pendants was observed to aggregate in solution, giving a FITC lifetime of 0.5 ns^15^; upon degradation of the oligomer by cathepsins the resulting FITC-lysine monomers reported a restored lifetime of 4 ns.

Aside from FITC, other small molecule fluorophores have reported faster lifetimes when associated with large polymers, aggregates, or biomacromolecules, than when measured as smaller entities^11, 16, 17^. While the reverse trend has also been reported^13, 18^, these findings suggest that fluorescence lifetime is sensitive to local aggregation. From this, we anticipated that incorporating FITC into a self-assembling structure would result in similar lifetime shifts which could serve as a signal for imaging biological events.

Reporter aggregation is a strategy proposed to improve the efficacy of small molecule imaging of biological events^19^. Aggregation of a non-targeted, small molecule reporter inhibits mobility and promotes retention at the site of activation^19–21^, providing better spatial resolution usually achieved with active-targeting or covalent-labeling imaging strategies. One such aggregation system is built upon the final step in the biosynthesis of luciferin^22^ and involves the condensation of the 1,2-aminothiol group of free cysteine and the nitrile group of 2-cyanobenzothiazole^23^ (CBT). Creating a molecule bearing both functionalities results in a compound that can condense into cyclic oligomers and subsequently self-assemble into nanostructures. This reaction is rapid^24^ (with a second-order rate constant of 9.19 M^−1^s^-1^) and can be controlled by caging portions of the 1,2-aminothiol group to create a bio-responsive aggregation system^23^. This system has been used as a platform for a wide variety of imaging methods^25–34^, including steady-state fluorescence imaging^20, 35–38^, but it has not been examined as a platform for time-resolved fluorescence imaging.

Considering the inherent sensitivity of fluorescence lifetime to aggregation, this triggered aggregation system may serve as a means of producing contrast in the lifetime of a covalently-linked fluorescent reporter. This scaffold could then be caged by substrates or other moieties to facilitate smart imaging in response to enzymes or cellular conditions of interest, a strategy thoroughly demonstrated by previous work^20, 25–33, 35–38^. For instance, it is reported that many cancer cell lines produce increased levels of glutathione (GSH) compared to their normal tissue counterparts^39–41^; caging the sulfhydryl group of the scaffold with a reduction-sensitive disulfide bond would facilitate glutathione-responsive lifetime imaging (Figure 1).

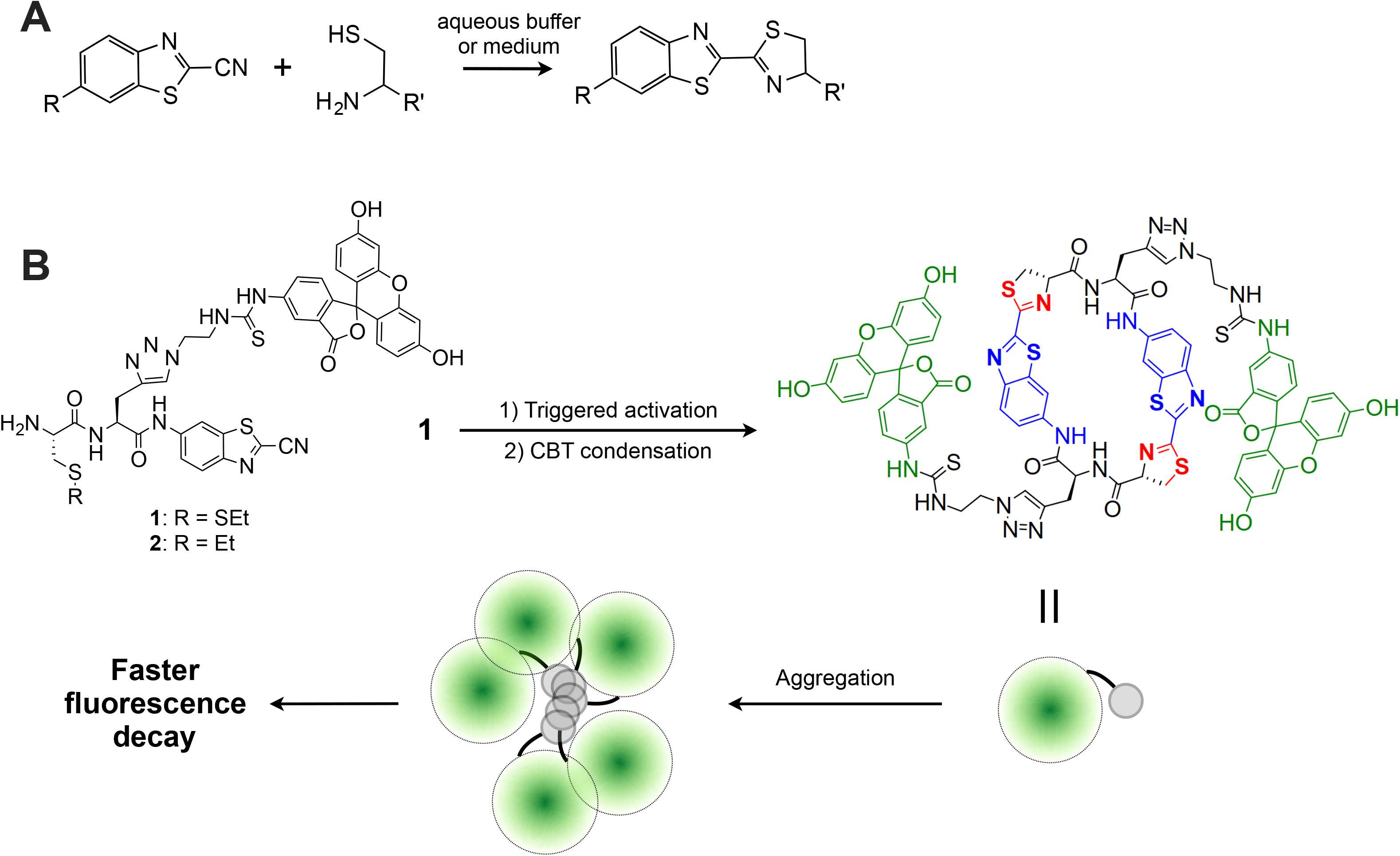

## Results and Discussion

### Characterization of GSH-triggered aggregation

We designed compound **1** by attaching FITC to a simple CBT scaffold (Scheme S1) and tested its response to GSH in PBS buffer. Compound 1 as a 100 μM,10 μM, or **1** μM solution in PBS (pH = 7.4) was incubated with 10 mM GSH, and aggregation of the activated (reduced) compound was detected by Nanoparticle Tracking Analysis (NTA). Nanoparticles of 150-200 nm average size were detected when **1** was incubated with GSH (Table S2), similar in size to other iterations of this scaffold^25, 33, 34, 37, 42^.

### Stead state electronic absorption and emission

To determine the influence of this aggregation on the nature of the attached FITC fluorophore, we examined the steady state electronic absorption, excitation, and emission spectra of **1** before and after activation (Figures 2 and 3). The absorption of the 100 μM activated solution of **1** (Figure 2A, black solid) shows two predominate inhomogeneously broaden peaks centered at 2.58 eV, and 3.29 eV. The absorption of activated **1** also shows a steady rise in absorptivity moving toward lower energy caused by scattering (Figure 2A). The broadening of the FITC absorption, 2.58 eV, could be resulting from a distribution of polymer sizes upon reaction which then perturbs the FITC moiety in a distribution of arrangements. The reaction, in theory, should not alter the FITC moiety but once the activated product of **1** is aggregated many different nano-environments may alter the FITC moiety to a large degree, resulting in broadening. The broadened peak centered at 3.29 eV is assigned to the product oligomer of **1.** Evidence for the assignment can be seen in Figure 3 as overlapping excitation spectra from 2.74 eV (453 nm) emission (yellow solid squares) with the absorption. This also gives more evidence for the new emission seen in Figure 2B, at 2.74 eV (453 nm) predominantly coming from the resulting product **1** oligomer. The excitation spectrum from the emission at 2.41 eV (515 nm) (Figure 3B, green dashed) is broadened compared to **1** alone (Figure 3A, green dashed), as well as having a new feature described as a “fat” tail continuing towards higher energy. In **1** alone the emission at 2.74 eV (453 nm) dominantly comes from the FITC moiety but in the 100 μM reaction of **1** seemingly everything contributes towards the emission. We believe this contributes to the broadening seen in the emission (Figure 2B).

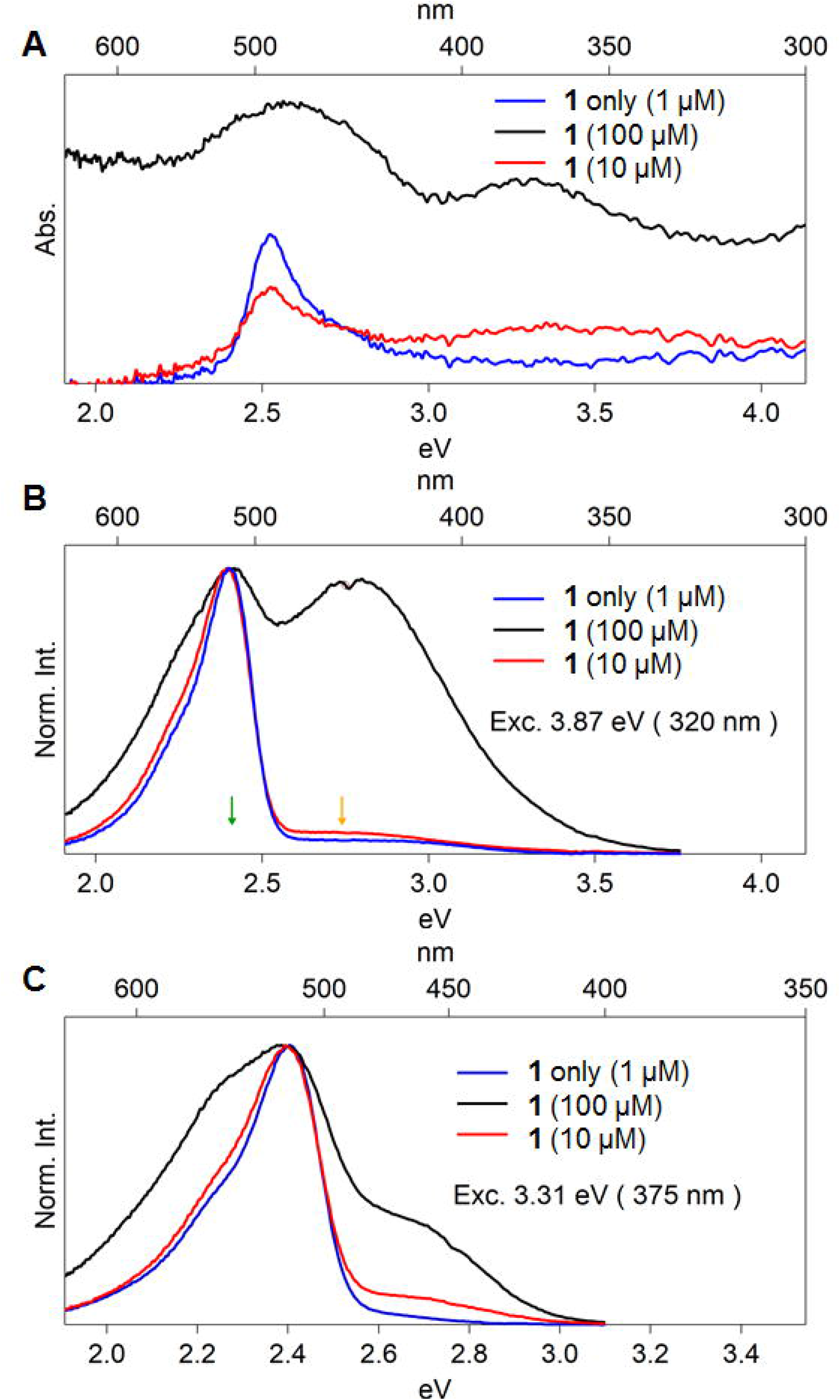

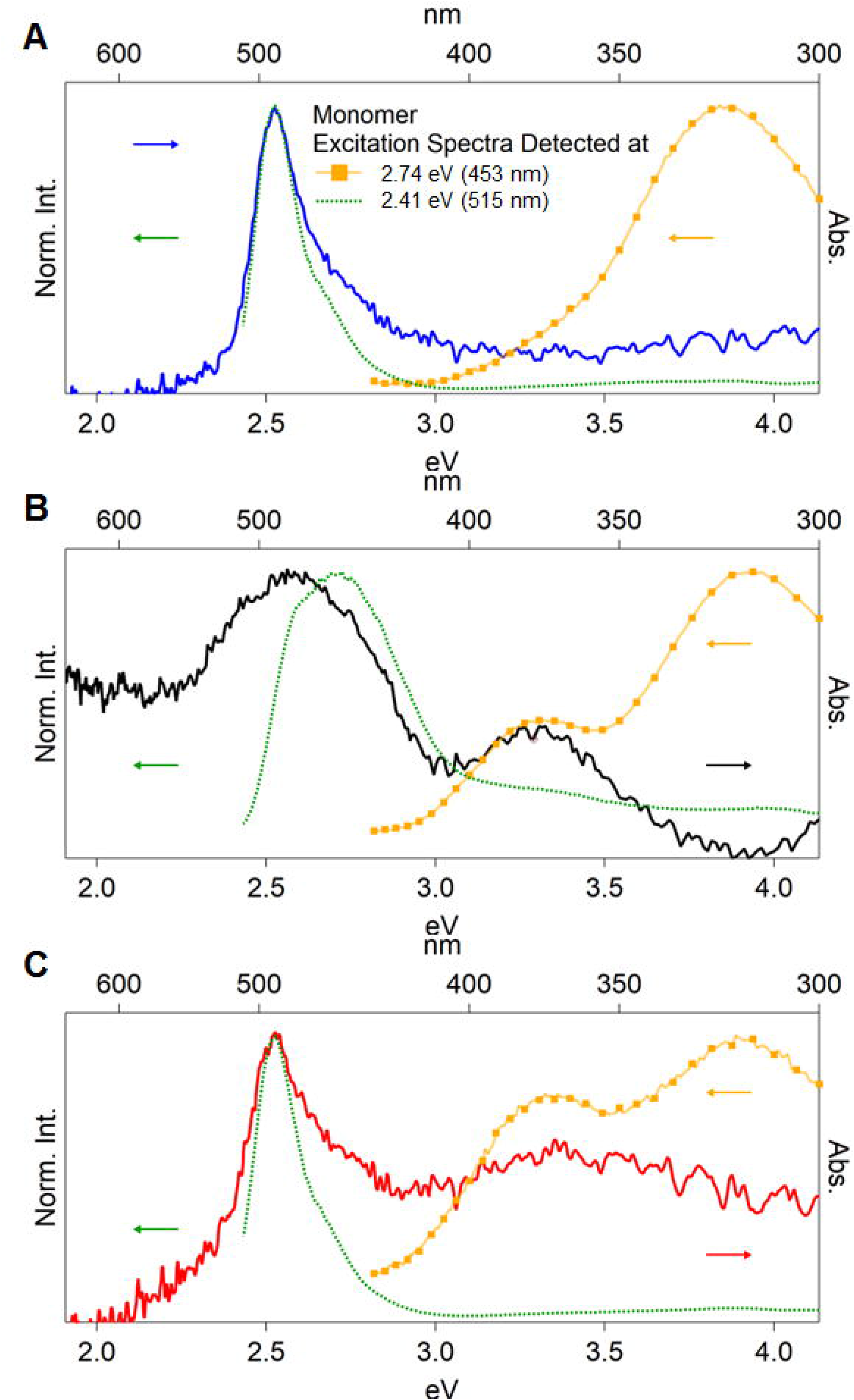

Emission from **1** alone has two assigned features: 2.40 eV (516 nm) from the FITC, and 2.92 eV (425 nm) from CBT (Figure 2B and Figure S2). Upon activation emission from the FITC broadens from a FWHM of 0.21 eV to at least 0.31 eV. A new feature emerges centered at 2.74 eV (453 nm) which is assigned as the emission from the resulting product. The lack of the 2.74 eV (453 nm) emission seen in **1** alone gives strong evidence for the identity (Figure 3A, yellow dotted).

### Steady state emission intensity and anisotropy

With evidence that **1** exhibits spectral changes upon reduction-triggered aggregation, we turned to examine other emission characteristics of the compound that could be influenced by aggregation. To determine if these changes were due to reduction-triggered aggregation of **1**, we also examined control compound **2**, which lacks the disulfide bond necessary to trigger activation and aggregation in response to GSH. Aggregation of fluorescein is known to alter its spectral properties, including emission lifetime^43^ and quantum yield^44^, which reduces emission intensity. Fluorescein is also known to undergo homo-FRET (Fluorescence Resonance Energy Transfer, taking place between two identical fluorophore molecules), causing a depolarization of fluorescence emission from the sample^43, 45^; this effect increases as more fluorophores come within FRET distance of each other^45^. We examined whether any of these trends were observed in **1** by measuring its steady-state emission intensity, anisotropy (polarization), and lifetime values at different concentrations, and comparing the results to that of **2** (Figure 4).

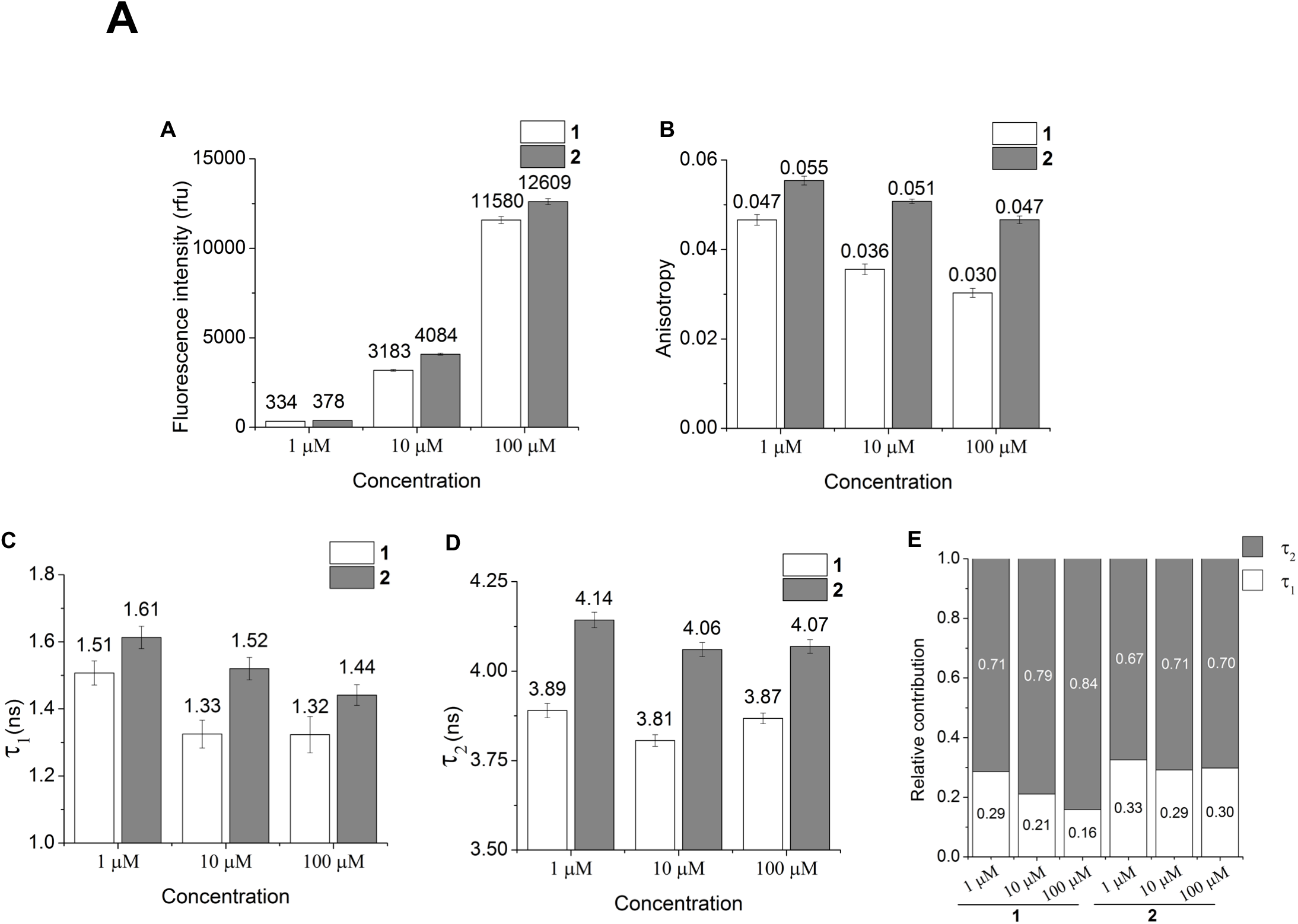

The emission intensity of both compounds was equal at 1 μM, while at higher concentrations a statistically significant reduction in emission intensity was measured for **1** when compared to **2** (*p* < 0.05), suggesting that triggered aggregation is reducing the quantum yield of FITC in **1** at these concentrations. At all concentrations, the anisotropy of **1** was lower than that of **2**; as concentration increased, the anisotropy decreased for both compounds. The consistently lower anisotropy of **1** is attributed to aggregation of the oligomer products, reasonably bringing multiple FITC molecules within homo-FRET distance of each other. Models of aggregated CBT systems suggest that these aggregates pack together with an intermolecular distance of 0.37 nm^46^, which is well within the measured 5.3-5.7 nm Förster radius of FITC^45^. We attribute the concentration-dependent reduction of anisotropy possibly to low solubility for both compounds in this concentration range; indeed, general insolubility would result in compound packing to some extent and thus facilitate homo-FRET and a resultant reduction in anisotropy. While a distinct and significant (*p* < 0.05) difference in anisotropy is observed between **1** and the inactive control **2**, the ideal probe should exhibit minimal self-depolarization before activation (while in the form of a single molecule, such as **2**), but exhibit dramatic depolarization upon activation, attributed exclusively to the aggregation process. While this data indicates that aggregated **1** has consistently lower anisotropy than inactive small molecule **2**, a cleaner, binary change in anisotropy is desired. Overall, the reduced emission intensity and anisotropy of **1** suggest that aggregation is indeed influencing the attached FITC fluorophore. We next investigated whether the aggregation process altered the emission lifetime of FITC to an extent that would enable lifetime-based imaging.

### Time Resolved Emission Spectra

Time Resolved Emission Spectra (TRES) scans were performed to obtain the lifetimes of fluorescence emission. For 12 nm steps starting at 400 nm and ending at 700 nm an emission lifetime was collected from a 375 nm laser using TCSPC. From the lifetime map the corresponding time resolved emission can be reconstructed after correcting for the detector’s response. The resulting TRES maps are shown in Figures S1, S3, and S4, and Table S1. Each lifetime from the TRES scan was fitted using the model shown in Table S1, using iterative reconvolution with the instrument response function (IRF), shown in Figure S1. Two chosen fits from the TRES map can be seen in Figures S1 (2.38 eV, 520 nm) and S3 (3.01 eV, 412 nm). Using this method, we can measure lifetimes down to ~200 ps.

To better understand the dynamics, we constructed a Transient Emission Normalized Area Spectroscopy (TRANES) scan where each trace’s area is normalized to unity^47^. This procedure reveals isoemissive points similar to isosbestic points in transient absorption spectroscopy (TAS). The resulting TRANES can be seen in Figure S5. First looking at the normal TRES for **1** alone, Figure S4a, the large peak at ~2.8 eV, rising from 1.52 ns then decaying after 3.03 ns is assigned to the CBT emission. The peak at 2.4 eV which appears at 1.52 ns reaching a maximum at 3.03 ns then decaying at >18.24 ns is assigned to the FITC emission. Upon the reaction, Figure S4b, the CBT emission is now the prominent peak and a new emission appears filling the void at 2.7 eV. The new emission at 2.7 eV is assigned to the resulting product material from the condensation. The FITC emission at 2.4 eV is also broaden compared to the monomer but has similar decay. Comparing the TRES map and TRANES map, Figure S4b and S5b respectfully, there is now a clear isoemissive point at 2.45 eV. The emergence of the isoemissive point upon reaction is evidence of two emitting species, one being the product material from the condensation and the other, most likely being unreacted starting material. The trend continues in the reaction at 10 μM where the isoemissive point is still present but not as clean as in the reaction at 100 μM and the decay line shape narrows with strong similarities to the monomer in both TRES and TRANES, (Figure S4c and S5c), thus giving more evidence there is both unreacted monomer and the product material in both concentrations of reactions.

Fluorescence emission lifetime at 2.38 eV (532 nm) emission (FITC region) was 3.96 ns (100%) for **1** only, consistent with the literature-reported lifetime of FITC small molecule^48, 49^ (Figure S1). Upon reaction with GSH, the lifetime of **1** changed to 3.89 ns (71.37%) for the 1 μM reaction mixture, 3.81 ns (78.91%) for the 10 μM reaction, and 3.87 ns (84.15%) for the 100 μM reaction (Figure 4D, E). In comparison, the lifetime of **2** was 4.14 ns (67.47%) at 1 μM, 4.06 ns (70.82%) at 10 μM, and 4.07 ns (70.20%) at 100 μM (Figure 4D, E). This major component is faster in **1** than in **2** at all concentrations tested, a trend also seen in the second, minor component of both compounds (Figure 4C). Overall, **1** mixed with a reducing agent exhibits a faster lifetime than both compound **2** and compound **1** in the absence of a reducing agent, suggesting that both the reduction-sensitive disulfide bond of **1** and the presence of a reducing agent are required to shorten the lifetime of the attached FITC.

Aside from the FITC region, we also found a dramatic lifetime change at 3.01 eV (412 nm, CBT region) (Figure S3, S4, and Table S1). The lifetimes from the fit are 0.32 ns (96.62%), 0.42 ns (95.81%), and 8.84 ns (98.04%), for **1** alone, activated **1** at 10 μM, and activated **1** at 100 μM, respectively. This may suggest the packing of the aggregates after CBT condensation relies more on the CBT moiety, while the FITC moieties are arranged more toward the peripheral region. This observed dramatic lifetime change in the CBT moiety is difficult to apply to lifetime-based imaging because of significant cellular autofluorescence in the blue region of its emission profile (3.01 eV, 412 nm).

### Fluorescence Lifetime Imaging Microscopy of mammalian cells

Altered fluorescence lifetime has been used as a readout for many biochemical assays performed at the cellular level^10^. As **1** demonstrates reduction-triggered contrast in lifetime, we considered if this lifetime change could still be detected in the complex environment of the cell. First, we determined cellular uptake of the compounds by incubating HeLa cells with compound **1** (pre-activated by incubation with GSH), or with compound **1** or **2** alone. We examined the uptake of the compounds using fluorescence microscopy and flow cytometry (Figure 5). Fluorescence microscopy (Figure 5A) revealed notable differences in uptake patterns between the compounds: the pre-activated aggregates of **1** demonstrated high but diffuse uptake; compound **1** showed slightly higher intensity than **2** in HeLa cells, but **1** appears to be localized to particular areas within the cells, while **2** demonstrates the diffuse staining seen for pre-activated **1**. The relative uptake in these samples was corroborated by flow cytometry results (Figure 5B, C).

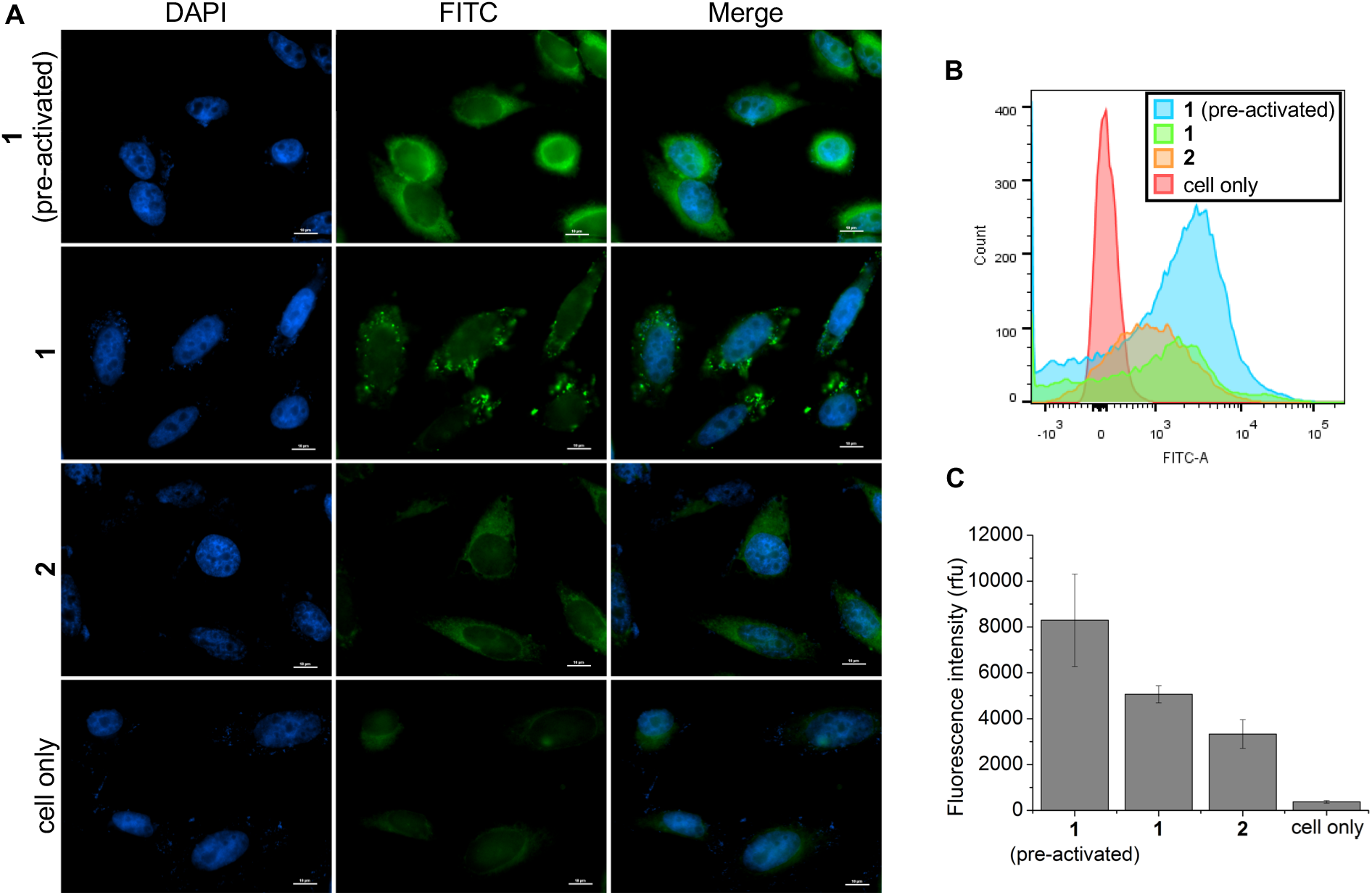

To determine if the lifetime change of **1** could be detected in cells, HeLa cells were incubated with probes and lifetimes were measured using Fluorescence Lifetime Imaging Microscopy (FLIM, Figure 6A). The measured lifetimes for all three samples were best fit using bi-exponential functions (Figure 6B, D), in agreement with our previous lifetime measurements (Figure 4C-E). In cells, compound **1** preincubated with GSH exhibited shorter values for both lifetime components when compared to **2** or **1** added alone. This is also in agreement with the previous trend, where **1** exposed to a reducing agent reported faster lifetime components than 2. It is notable than in these cell samples, the faster of the two lifetime components (τ_2_) became the major component for pre-activated **1** compared to **1** or **2** alone, resulting in a dramatic decrease (~ 0.9 ns) in the average lifetime of the reporter (Figure 6C, D). It is of note that when compound **1** was added to HeLa cells without addition of an exogenous reducing agent, no reduction in lifetime was observed in comparison with 2. This may suggest that aggregation is not achieved to a suitable extent in this context, for example by condensation of the reduced CBT group onto thiol-containing biological molecules^50–52^ instead of another molecule of **1**. Judging from the greater uptake of **1** into cells over **2** (Figure 5C), it is possible that **1** is activated (reduced) in cells but is interrupted before condensing with another molecule of **1**, resulting in greater uptake and retention in cells when compared to **2**, but without lifetime change that may come with aggregation. A more efficient self-assembly system may be achieved by tailoring the CBT scaffold reactivity^37, 42^.

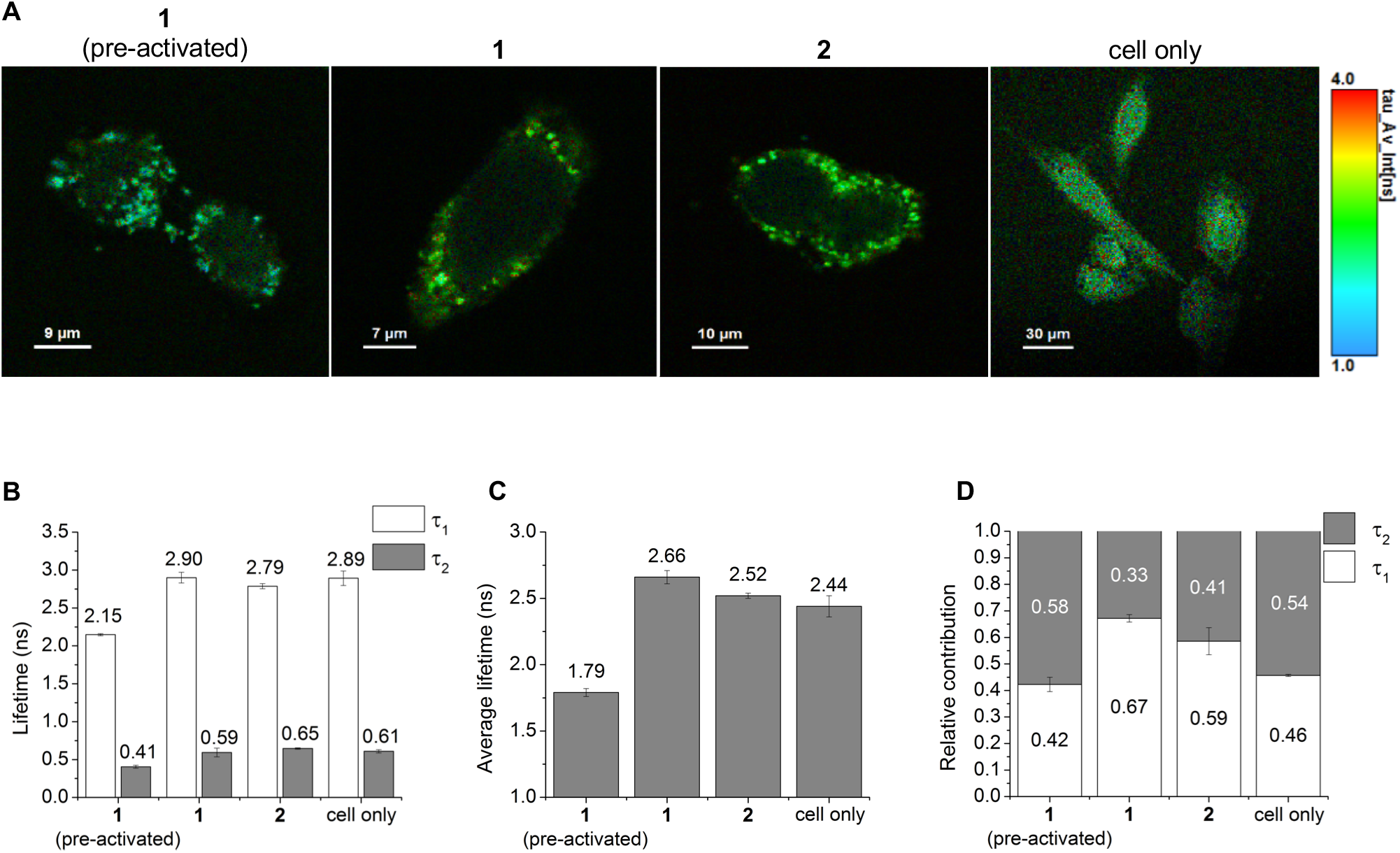

## Conclusions

In conclusion, we report a reduction-sensitive fluorescent probe that exhibits an altered emission lifetime upon exposure to reducing conditions. A decrease in lifetime is observed when **1** is measured in the excitation and emission ranges of FITC, suggesting that the fluorophore is indeed influenced by the aggregation of the CBT scaffold. From this data, several improvements should be considered. In solution, notable lifetime changes are observed in the emission range of FITC only when compound **1** is incubated with a reducing agent such as GSH; the presence of a reduction-sensitive disulfide bond and a reducing agent seem to be required to facilitate the lifetime shift. However, it is important to discern the underlying mechanism(s) influencing the lifetime during aggregation: fluorophore aggregation can be associated with various quenching processes^14, 53^, but in fluorescein and other reporters only some of these processes alter the emission lifetime^43^. With a better mechanistic understanding, the fluorophore and CBT scaffold could be tailored to direct these quenching processes and create a more dramatic lifetime change upon activation. While detected using FLIM, the lifetime decrease of **1** was not generated *in situ* in cells, suggesting that a careful selection of fluorophore and scaffold may also be required to allow the aggregation process to occur in the complex environment found within the cell. Along with this, the influence of concentration on the polarization and lifetime values leads us to believe a ‘solubility switch’ structure should be considered, in which the reduction reaction removes a highly water-soluble functional group, improving the solubility of the monomer while retaining the aggregation potential of the reduction product. Further study of this type of aggregation-induced lifetime contrast system can elucidate the mechanisms contributing to the observed lifetime change and illuminate synthetic design strategies to enhance the lifetime contrast produced by probe activation.

## Experimental Methods

### Chemical Synthesis

Experimental procedures and characterization data are provided in the Supplementary Information.

### Steady state electronic absorption and emission, TRES

A solution of **1** (200 uM) in PBS (pH 7.4) was incubated with GSH (10 mM) and NaHCO_3_ (40 mM) and stirred for 30 min to **1** h. Aliquots of the reaction mixture were then sequentially diluted using a 100 μM dilution factor and a 10 μM dilution factor. All solutions were in PBS buffer at pH 7.4. It is important to note spectra were measured *in vitro* as no purification or work up was done on the final reaction mixture before acquiring. Measuring in this manner best simulates a similar inhomogeneous mixture found in the triggered system. Steady state electronic absorption spectra were collected on a UV-2550 Shimadzu. Steady state emission, excitation, synchronous scan, and time resolved emission spectra (TRES) were collected on an Edinburgh Instruments FLS1000 Spectrometer. All spectra were collected with 1 cm path length quartz cuvette unless otherwise stated. Spectra was collected on a 1 μM solution of **1** alone for comparison.

### Steady state fluorescence intensity and anisotropy, fluorescence lifetime

To **1** or **2** was added PBS buffer (1x, pH = 7.4) to make 90% volume of the final solution at 1.11x the desired final concentration, then the solution was vortexed for 10 sec. A separate PBS solution of GSH (100 mM) and NaHCO_3_ (400 mM) was prepared, and the resulting solution was filtered with a 0.2 micron syringe filter. The filtered solution was diluted 10-fold into the compound-PBS solution, to produce a final solution at 100% volume, containing 10 mM GSH, 40 mM NaHCO_3_, and a compound concentration of 100 μM, 10 μM, or 1 μM. The solution was vortexed again for 10 sec to mix, then allowed to sit at room temperature for 30 min to 1 hour before measurements were collected. Fluorescence lifetime measurements were collected on an Edinburgh Instruments FLS1000 Spectrometer, excitation 488 nm, emission 520 nm. Samples for emission intensity and anisotropy were prepared similarly in a black-bottom 96-well plate for fluorescence measurements. Fluorescence intensity and anisotropy was measured on a SpectraMax M5 plate reader, with an excitation wavelength of 488 nm, and an emission wavelength of 518 nm.

### Nanoparticle tracking analysis

To **1** or **2** was added PBS buffer (1x, pH = 7.4) to make 90% volume of the final solution at 1.11x the desired final concentration, then the solution was vortexed for 10 sec. A separate PBS solution of GSH (100 mM) and NaHCO_3_ (400 mM) was prepared, and the resulting solution was filtered with a 0.2 micron syringe filter. The filtered solution was diluted 10-fold into the compound-PBS solution, to produce a final solution at 100% volume, containing 10 mM GSH, 40 mM NaHCO_3_, and a compound concentration of 100 μM, 10 μM, or 1 μM. The solution was vortexed again for 10 sec to mix, then allowed to sit at room temperature for 30 min to 1 hour before measurements were collected. NTA was measured using a NanoSight NS300. To retain the correlation of size to compound concentration, the solutions were injected without further dilution. Sample flow rate was set to 50 μL/min, and data was acquired as 5 replicates of 60 sec captures. Data was collected using a Blue488 laser, and all measurements were performed at 25 C as maintained by the instrument. Particle size and particle concentration were calculated in accordance with the software default settings.

### Cell culture and treatment

HeLa cells were cultured at 37□°C in 10□cm dishes containing Dulbecco’s Modified Eagle’s medium (DMEM) supplemented with 10% fetal bovine serum and antibiotics (100□U/mL penicillin, 100□μg/mL streptomycin) under 5% CO_2_ and 95% humidity.

### Flow cytometry

HeLa cells were seeded in a 24-well plate at a density of 3□×□10^5^ cells/well (total of 12 wells) and cultured overnight. The next day, cells were incubated with pre-activated **1** (2 μM, 3 wells), **1** only (2 μM, 3 wells), **2** only (2 μM, 3 wells), or no probe (3 wells) for 1 hour, then washed with PBS buffer and digested with 0.25%trypsin. Cells were collected into Eppendorf tubes, washed with PBS buffer 3 times at 1250□×□*g* for 3min, and resuspended in 200□μL PBS buffer. The fluorescence of cell samples (3 samples for each data point) was analyzed with a flow cytometer (Canto II, Becton Dickinson and Company, USA).

### Fluorescence cell staining

HeLa cells were seeded on glass coverslips (0.13-0.16 mm thickness) at a density of 3 × 10^4^ cells/well (total 4 wells) and cultured overnight. The next day, cells were incubated with pre-activated **1** (2 μM, 1 well), **1** alone (2 μM, 1 well), **2** alone (2 μM, 1 well) or no probe (1 well) for 1 hour, then fixed with 4% paraformaldehyde at 37 °C for 10 min. The samples were washed with PBS 3 times, 5 min each. Cells were then incubated with DAPI for 5 min at room temperature. Coverslips were mounted on slides using ProLong Gold Antifade Mounting media and were used for fluorescence microscopy (Nikon Ti2, Japan) and fluorescence lifetime imaging microscopy described below.

### Fluorescence lifetime imaging microscopy

Lifetime imaging was performed on a Leica TCS-SP8 microscope (Leica Microsystems GmbH, Wetzlar, Germany) with a PicoHarp 300 (PicoQuant, Berlin, Germany) lifetime module, possessing a resolution of 4 ps. Excitation was generated by a 488 nm white light laser with a 40 Hz pulse for 5-10 cycles. Emission was centered at 516 nm (range of 512 to 520 nm). Images were processed with SymPhoTime 64 software (PicoQuant, Berlin, Germany). Results were processed with a multi-exponential (“*n*-exponential” function) tail fit, increasing the number of components until the resulting function provided a chi-squared value of approximately 1.0.

## Acknowledgements

This work was supported by research grants to Prof. L. Cui from the University of New Mexico (UNM Startup Award), the National Institute of General Medical Sciences of National Institutes of Health (Maximizing Investigators’ Research Award for Early Stage Investigators, R35GM124963), and the Career Development Award from the Department of Defense (W81XWH-17-1-0529). We are grateful to the support from the Department of Chemistry and Chemical Biology, University of New Mexico (UNM), the UNM Comprehensive Cancer Center and the National Cancer Institute of the United States (P30CA118100). NMR spectra were collected in part from the NMR Facility, Department of Chemistry and Chemical Biology, UNM, and from the Department of Medicinal Chemistry, College of Pharmacy, University of Florida (UF). Mass spectrometry services were provided in part by the Mass Spectrometry Facility, Department of Chemistry and Chemical Biology, UNM, and from the Mass Spectrometry Research and Education Center, Department of Chemistry, UF (NIH S10 OD021758-01A1). Nanoparticle Tracking Analysis instrumentation was provided by the UF Interdisciplinary Center for Biotechnology Research. Images in this paper were generated in the UNM Comprehensive Cancer Center Fluorescence Microscopy Shared Resource.

## Supporting Information

The Supporting Information is available on the bioRxiv preprint site.

