## Supplementary information for "Responsive fluorophore aggregation provides spectral contrast for fluorescence lifetime imaging"

### 1. General Reagents and Instruments

Solvents and chemicals were purchased from commercial sources and used directly without further purification. Flash chromatography was performed manually with Agela Technologies Flash Silica (40-60 µm, 60 Angstroms). HPLC was performed on a Dionex UltiMate 3000 with a pump and an in-line Diode Array Detector (DAD-3000). A reverse-phase C18 (Teledyne, 5 µm, 10 x 250 mm) column was used for analysis and semi-preparation. $^{1}$H NMR spectra were recorded using either a Bruker Avance III 300, Bruker Avance 500, or a Bruker Avance Neo 600, and chemical shifts were reported in ppm with either TMS or deuterated solvents as internal standards (TMS, 0.00; $CDCl_3$, 7.26; MeOD, 3.31). $^{13}$C NMR spectra were recorded at 75.4 or 125.7 MHz, and chemical shifts were reported in ppm with deuterated solvents as internal standards ($CDCl_3$, 77.0; MeOD, 49.15). Nanoparticle Tracking Analysis (NTA) was performed on a NanoSight NS300. Steady state electronic absorption spectra were collected on a UV-2550 Shimadzu. Steady state emission, excitation, synchronous scan, and time resolved emission spectra (TRES) were collected on an Edinburgh Instruments FLS1000 Spectrometer. All spectra were collected with 1 cm path length quartz cuvette unless otherwise stated. Steady-state emission intensity and polarization were collected on SpectraMax M5 plate reader.

#### 2. Experimental Section

**Chemical Synthesis:** Experimental procedures and characterization data are provided in the Supplementary Information.

**Measurements of the optical properties:** Aliquots of the reaction mixture were taken at then diluted to a 100 µM dilution factor, then sequential dilutions down to 10 µM dilution factor. All solutions were in PBS buffer at pH 7.4. It is important to note spectra were measured *in vitro* as no purification or work up was done on the final reaction mixture before acquiring. Measuring in this manner best simulates a similar inhomogeneous mixture found in the triggered system. Steady state electronic absorption spectra were collected on a UV-2550 Shimadzu. Steady state emission, excitation, synchronous scan, and time resolved emission spectra (TRES) were collected on an Edinburgh Instruments FLS1000 Spectrometer. All spectra were collected with 1 cm path length quartz cuvette unless otherwise stated. Spectra was collected on a 1 µM of just the monomer for comparison.

**Steady state fluorescence intensity and anisotropy, and fluorescence lifetime**: To **1** or **2** was added PBS buffer (1x, pH = 7.4) to make 90% volume of the final solution at 1.11x the desired final concentration, then the solution was vortexed for 10 sec. A separate PBS solution of GSH (100 mM) and $NaHCO_3$ (400 mM) was prepared, and the resulting solution was filtered with a 0.2 micron syringe filter. The filtered solution was diluted 10-fold into the compound-PBS solution, to produce a final solution at 100% volume, containing 10 mM GSH, 40 mM $NaHCO_3$, and a compound concentration of 100 µM, 10 µM, or 1 µM. The solution was vortexed again for 10 sec to mix, then allowed to sit at room temperature for 30 min to 1 hour before measurements were collected. Fluorescence lifetime measurements were collected on an Edinburgh Instruments FLS1000 Spectrometer, excitation 488 nm, emission 520 nm. Samples for emission intensity and anisotropy were prepared similarly in a black-bottom 96-well plate for fluorescence measurements. Fluorescence intensity and anisotropy was measured on a SpectraMax M5 plate reader, with an excitation wavelength of 488 nm, and an emission wavelength of 518 nm.

**Nanoparticle tracking analysis:** NTA was measured using a NanoSight NS300. To retain the correlation of size to compound concentration, the solutions were injected without further dilution. Sample flow rate was set to 50 µL/min, and data was acquired as 5 replicates of 60 sec captures. Data was collected using a Blue488 laser, and all measurements were performed at 25 C as maintained by the instrument). Particle size and particle concentration were calculated in accordance with the software default settings.

**Cell culture and treatment**: HeLa cells were cultured at 37 °C in 10 cm dishes containing Dulbecco's Modified Eagle's medium (DMEM) supplemented with 10% fetal bovine serum and antibiotics (100 U/mL penicillin, 100 µg/mL streptomycin) under 5% $CO_2$ and 95% humidity.

**Flow cytometry**: HeLa cells were seeded in a 24-well plate at a density of $3 \times 10^5$ cells/well (total of 12 wells) and cultured overnight. The next day, cells were incubated with pre-activated **1** (2 µM, 3 wells), **1** only (2 µM, 3 wells), **2** only (2 µM, 3 wells), or no probe (3 wells) for 1 hour, then washed with PBS buffer and digested with 0.25% trypsin. Cells were collected into Eppendorf tubes, washed with PBS buffer 3 times at 1250 × *g* for 3min, and resuspended in 200 µL PBS buffer. The fluorescence of cell samples (3 samples for each data point) was analyzed with a flow cytometer (Canto Ⅱ, Becton Dickinson and Company, USA).

**Fluorescence cell staining**: HeLa cells were seeded on glass coverslips (0.13-0.16 mm thickness) at a density of $3 \times 10^4$ cells/well (total 4 wells) and cultured overnight. The next day, cells were incubated with pre-activated **1** (2 µM, 1 well), **1** alone (2 µM, 1 well), **2** alone (2 µM, 1 well) or no probe (1 well) for 1 hour, then fixed with 4% paraformaldehyde at 37 °C for 10 min. The samples were washed with PBS 3 times, 5 min each. Cells were then incubated with DAPI for 5 min at room temperature. Coverslips were mounted on slides using ProLong Gold Antifade Mounting media and were used for fluorescence microscopy (Nikon Ti2, Japan) and fluorescence lifetime imaging microscopy described below.

**Fluorescence lifetime imaging microscopy:** Lifetime imaging was performed on a Leica TCS-SP8 microscope (Leica Microsystems GmbH, Wetzlar, Germany) with a PicoHarp 300 (PicoQuant, Berlin, Germany) lifetime module, possessing a resolution of 4 ps. Excitation was generated by a 488 nm white light laser with a 40 Hz pulse for 5-10 cycles. Emission was centered at 516 nm (range of 512 to 520 nm). Images were processed with SymPhoTime 64 software (PicoQuant, Berlin, Germany). Results were processed with a multi-exponential ("*n*-exponential" function) tail fit, increasing the number of components until the resulting function provided a chi-squared value of approximately 1.0.

#### 3. Supplementary spectral data (Figures S1-S5, Tables S1-S2)

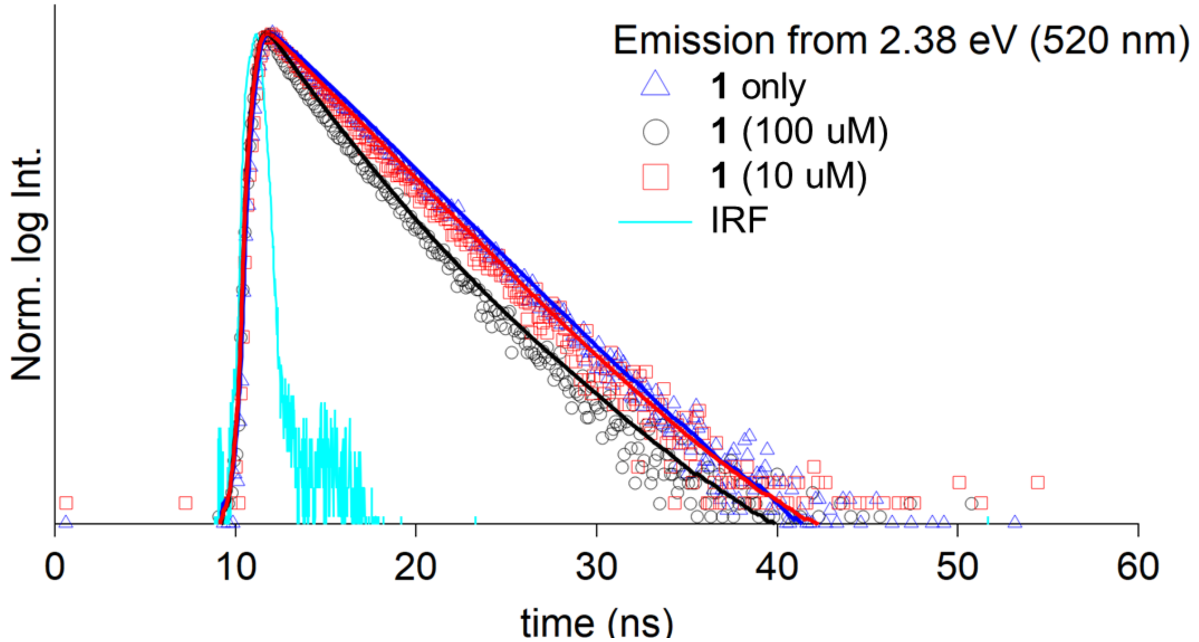

**Figure S1.** Fluorescence emission lifetime at 2.38 eV (530 nm) for **1** alone at 1 µM (blue triangles), or **1** at 100 µM reaction (black circles) or 10 µM (red squares) incubated with TCEP at pH 7.4. Fits to each trace are shown in solid. All kinetic fitting along with the model used can be seen in Supporting Information (Table S1).

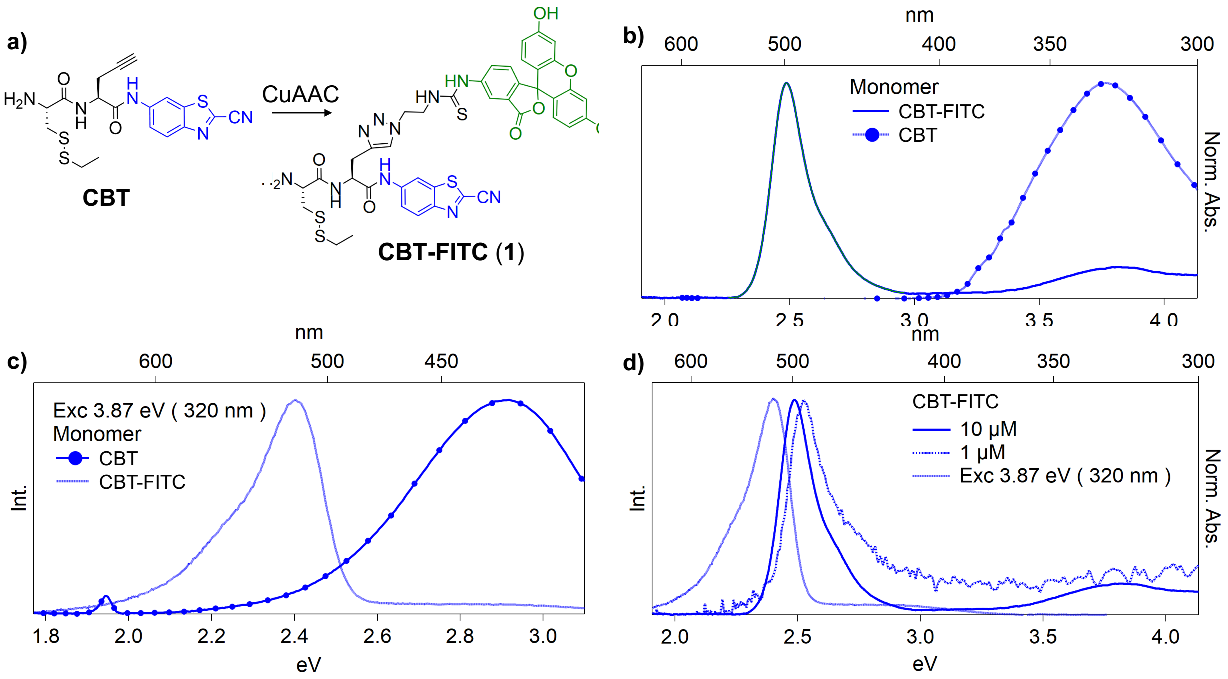

**Figure S2**. Steady state electronic absorption spectra of monomer. a) The chemical structure of the monomer CBT and CBT-FITC (compound **1**). b) Electronic absorption of the monomer with and without FITC, solid blue and dots, respectfully. The FITC contribution to the absorption is highlighted in green. c) The emission, upon 3.87 eV (320 nm) excitation, of both monomer with and without FITC, dashed and dots respectively. d) Electronic absorption of **1** at 10 μM (solid), at 1 μM (dashed), and emission from 3.87 eV (320 nm) excitation of the 1 μM concentration (dotted). All measurements were done at pH 7.4 in a PBS buffer solution.

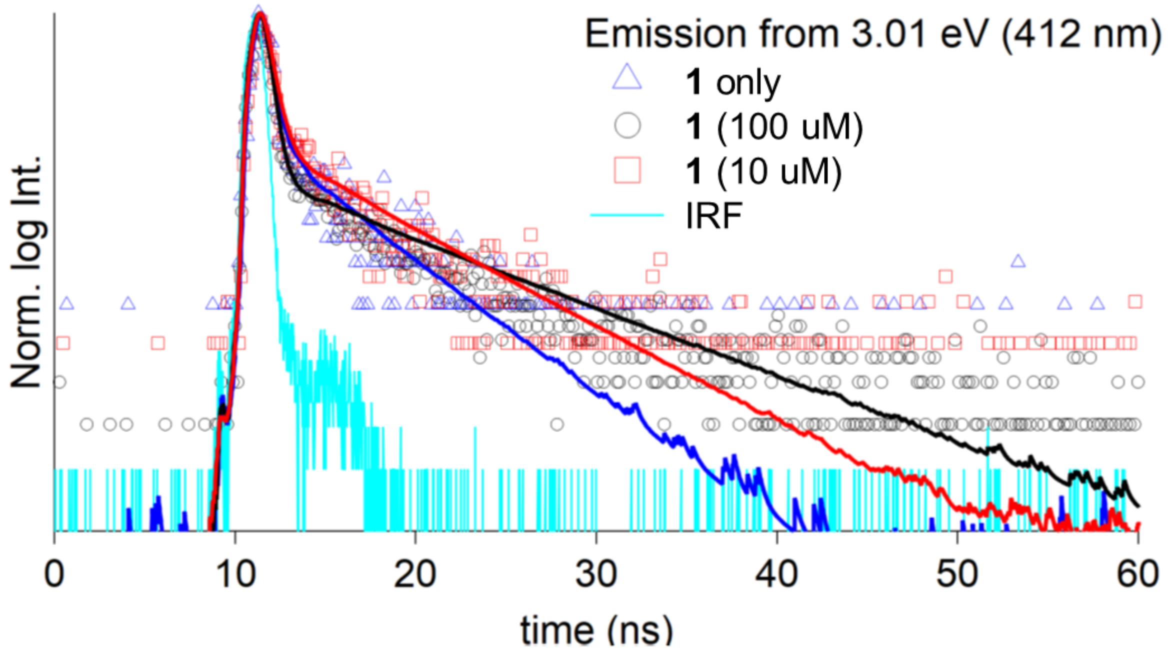

**Figure S3**. Fluorescence emission lifetime at 3.01 eV (412 nm) **1** alone at 1 µM (blue triangles), or **1** at 100 µM (black circles) or 10 µM (red squares) incubated with TCEP at pH 7.4. Fits to each trace are shown in solid.

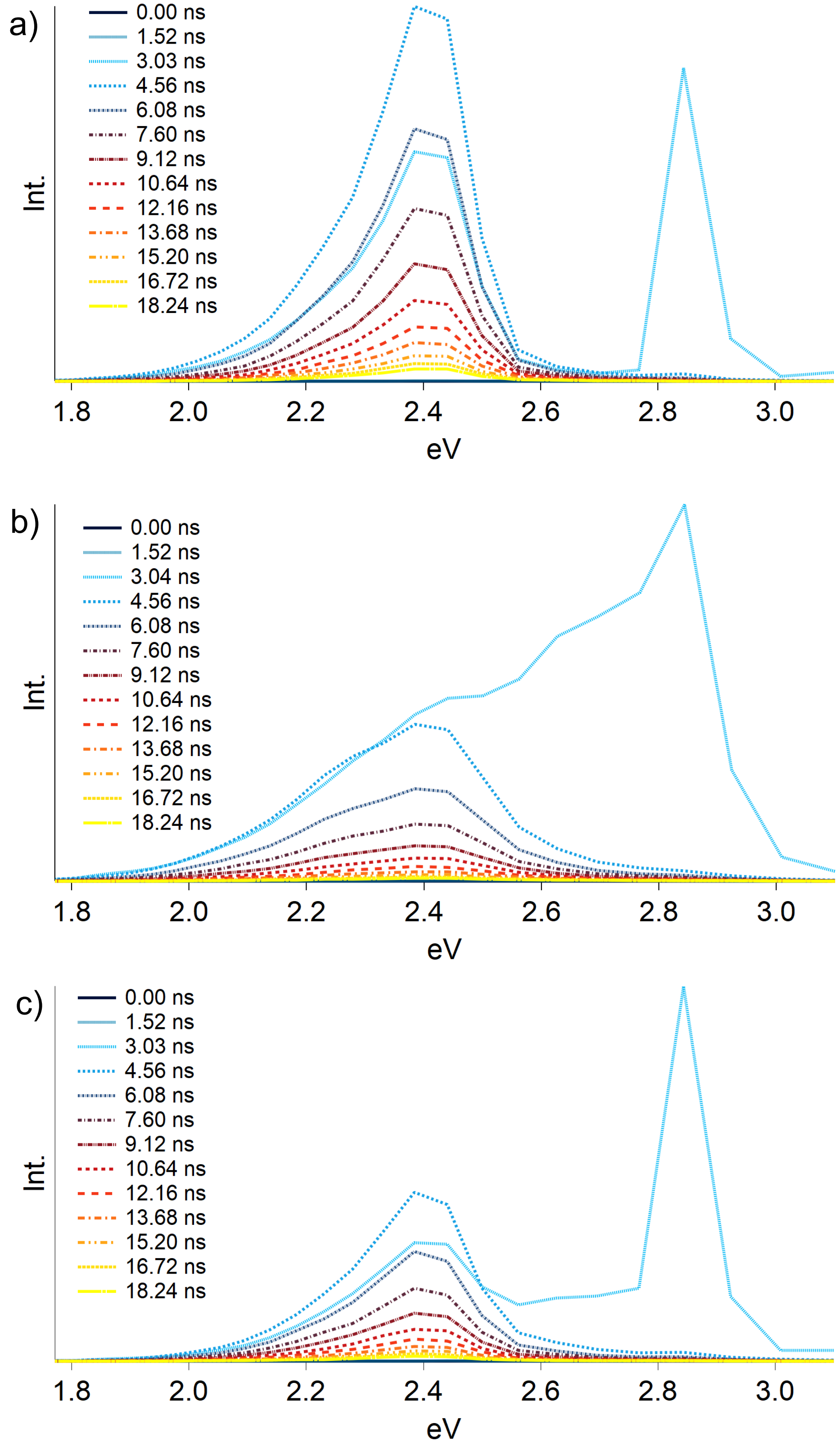

**Figure S4**. Time resolved emission spectroscopy scans for **1** alone at 1 µM (panel a), **1** at 100 µM incubated with TCEP (panel b), and **1** at 10 µM incubated with TCEP (panel c).

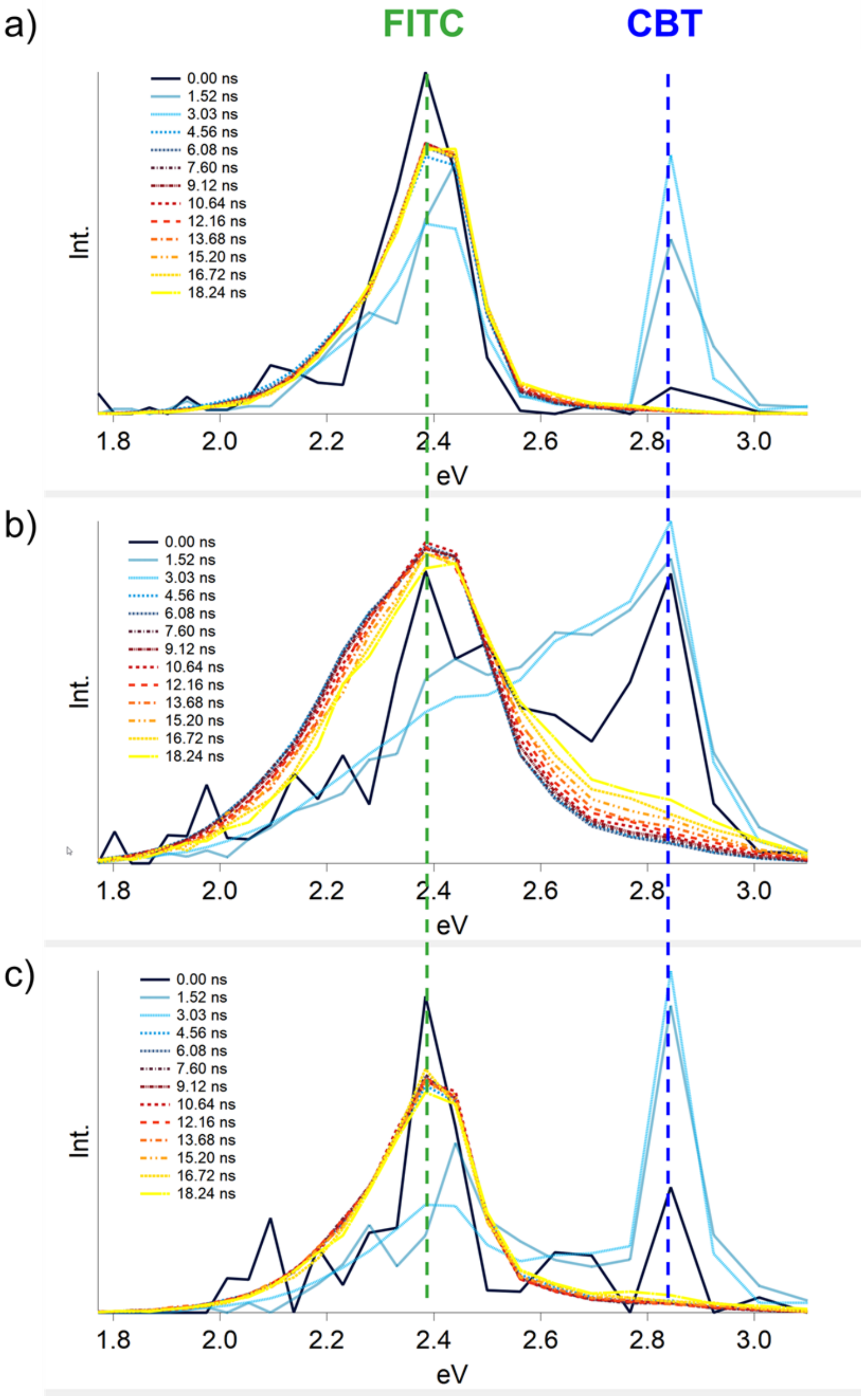

**Figure S5.** Transient Emission Normalized Area Spectroscopy (TRANES) scan of the traces shown in Figure S4. (a) **1** alone at 1 µM (blue triangles) and (b) **1** at 100 µM (black circles) or (c) 10 µM (red squares) incubated with TCEP at pH 7.4. Colored text labels (FITC, CBT) and corresponding dashed lines indicate peak assignments to portions of compound **1**.

| | $\tau_1$/ns ( $A_1$/% ) | | | $\tau_2$/ns ( $A_2$/% ) | | | $\tau_3$/ns ( $A_3$/%) | | |
|---|---|---|---|---|---|---|---|---|---|
| **eV ( nm )** | **Monomer 1 µM** | **Reaction 100 µM** | **Reaction 10 µM** | **Monomer 1 µM** | **Reaction 100 µM** | **Reaction 10 µM** | **Monomer 1 µM** | **Reaction 100 µM** | **Reaction 10 µM** |
| *3.09 (400)* | 0.19 (99.29) | 0.33 (97.89) | 0.32 (98.41) | 4.65 (0.70) | 7.96 (2.10) | 5.58 (1.58) | | | |
| *3.00 (412)* | 0.32 (96.62) | 8.84 (98.04) | 0.42 (95.81) | 4.53 (3.37) | 0.33 (1.95) | 6.23 (4.18) | | | |
| *2.92 (424)* | 0.01 (99.89) | 0.25 (99.28) | 0.15 (99.23) | 5.00 (0.10) | 8.05 (0.71) | 5.46 (0.76) | | | |
| *2.84 (436)* | 0.18 (99.93) | 0.28 (99.69) | 0.26 (99.8) | 8.40 (0.06) | 8.38 (0.30) | 7.04 (0.19) | | | |
| *2.76 (448)* | 0.47 (83.16) | 0.29 (99.22) | 0.34 (97.37) | 5.57 (16.83) | 6.61 (0.77) | 5.50 (2.62) | | | |
| *2.69 (460)* | 1.23 (56.87) | 0.32 (98.4) | 0.39 (94.85) | 5.77 (43.12) | 5.37 (1.59) | 4.72 (5.14) | | | |
| *2.62 (472)* | 1.93 (50.45) | 0.37 (96.06) | 0.47 (89.28) | 5.80 (49.54) | 4.77 (3.93) | 4.45 (10.71) | | | |
| *2.56 (484)* | 2.64 (53.91) | 0.46 (88.9) | 0.54 (71.9) | 5.57 (46.08) | 4.24 (11.10) | 3.44 (26.39) | | | 9.44 (1.70) |
| *2.49 (496)* | 3.74 (94.08) | 0.63 (61.65) | 2.01 (57.18) | 6.68 (5.92) | 3.26 (37.10) | 4.59 (42.81) | | 8.77 (1.25) | |
| *2.44 (508)* | 3.96 (100) | 3.32 (58.98) | 3.63 (96.43) | | 1.24 (39.42) | 7.05 (3.56) | | 8.29 (1.59) | |
| *2.38 (520)* | 3.96 (100) | 2.44 (82.02) | 3.79 (98.94) | | 4.96 (17.98) | 8.63 (1.05) | | | |
| *2.33 (532)* | 3.93 (100) | 2.60 (86.03) | 3.74 (98.28) | | 5.17 (13.97) | 8.04 (1.71) | | | |
| *2.27 (544)* | 3.86 (100) | 2.77 (92.06) | 3.63 (96.45) | | 5.64 (7.93) | 6.95 (3.54) | | | |
| *2.22 (556)* | 3.79 (100) | 2.86 (95.55) | 3.69 (98.6) | | 6.26 (4.44) | 8.56 (1.39) | | | |
| *2.18 (568)* | 3.70 (100) | 2.81 (94.8) | 3.64 (97.92) | | 5.96 (5.19) | 8.12 (2.07) | | | |
| *2.13 (580)* | 3.59 (100) | 2.85 (96.01) | 3.57 (98.2) | | 6.10 (3.99) | 9.05 (1.79) | | | |
| *2.09 (592)* | 3.44 (100) | 2.87 (97.72) | 3.52 (97.92) | | 6.97 (2.27) | 8.80 (2.08) | | | |
| *2.05 (604)* | 3.33 (100) | 2.79 (96.56) | 3.29 (93.79) | | 6.25 (3.43) | 6.80 (6.20) | | | |
| *2.01 (616)* | 3.20 (100) | 2.78 (96.56) | 3.24 (96.45) | | 6.10 (3.43) | 8.18 (3.54) | | | |
| *1.97 (628)* | 3.12 (100) | 2.79 (97.77) | 3.33 (97.07) | | 6.88 (2.22) | 9.21 (2.93) | | | |
| *1.93 (640)* | 3.00 (100) | 2.10 (57.28) | 2.29 (60.93) | | 3.60 (42.72) | 4.44 (39.06) | | | |
| *1.90 (652)* | 3.03 (100) | 2.05 (58.01) | 2.10 (64.54) | | 3.58 (41.98) | 4.67 (35.45) | | | |
| *1.86 (664)* | 2.83 (100) | 3.21 (60.32) | 1.24 (52.16) | | 1.21 (39.68) | 3.99 (47.83) | | | |
| *1.83 (676)* | 2.83 (100) | 3.30 (53.51) | 1.28 (50.07) | | 1.71 (46.48) | 3.95 (49.92) | | | |
| *1.80 (688)* | 2.63 (100) | 2.91 (100) | 3.22 (100) | | | | | | |
| *1.77 (700)* | 2.58 (100) | 2.88 (100) | 3.05 (100) | | | | | | |

$$f(t) = A_1 e^{-\tau_1 t} + A_2 e^{-\tau_2 t} + A_3 e^{-\tau_3 t}$$

**Table S1.** Resulting fits using reconvolution with IRF for each lifetime in the TRES scan. The kinetic model is shown above. Where there is no lifetime shown the model was reduced to the respective single or double exponential fit.

| Compound | 1 | | | 2 | | |
|---|---|---|---|---|---|---|
| **Concentration (uM)** | **1**[a] | **10** | **100**[b] | **1**[a] | **10** | **100**[b] |
| mean | 204 | 165 | 179 | 134 | 156 | 176 |
| mode | 147 | 120 | 137 | 102 | 113 | 173 |
| SD | 101 | 63 | 120 | 54 | 67 | 63 |
| particle conc. (particles/mL) | 8.80E+06 | 7.10E+08 | 1.13E+09 | 2.01E+07 | 2.44E+08 | 1.47E+09 |
| conc. SD | 1.33E+06 | 1.94E+07 | 7.48E+07 | 2.55E+06 | 7.24E+06 | 1.06E+07 |
| particles per frame (20-100 desired) | 0.4 | 45.4 | 60.2 | 1.2 | 13.3 | 53.2 |
| Noise | No | No | High | No | No | Detected |
| Validity of concentration measurement | Unreliable | OK | Unreliable | Unreliable | OK | Be cautious |

[a]The sample did not meet the ideal particles-per-frame threshold for accurate analysis, making the concentration data unreliable.

[b]The sample reported significant noise, making the concentration data uncertain or unreliable.

| Supplementary Movie | Compound | Concentration (µM) |
|---|---|---|
| S1 | **1** | 1 |
| S2 | **1** | 10 |
| S3 | **1** | 100 |
| S4 | **2** | 1 |
| S5 | **2** | 10 |
| S6 | **2** | 100 |

**Table S2.** Nanoparticle tracking analysis (NTA) measurements of particle size for **1** and **2** at 1 µM, 10 µM, and 100 µM, incubated with GSH (10 mM) at pH 7.4. Values were acquired from software calculations. Note: robust measurements require 20-100 sample particles in the measurement frame simultaneously throughout the measurement capture time. Excerpts of 10 sec from each video capture are provided as **Supplementary Movies S1-S6**, in the other Supporting Information files.

### 4. Chemical Synthesis and Characterization

Compounds used in the study were synthesized by the route outlined in **Scheme S1**.

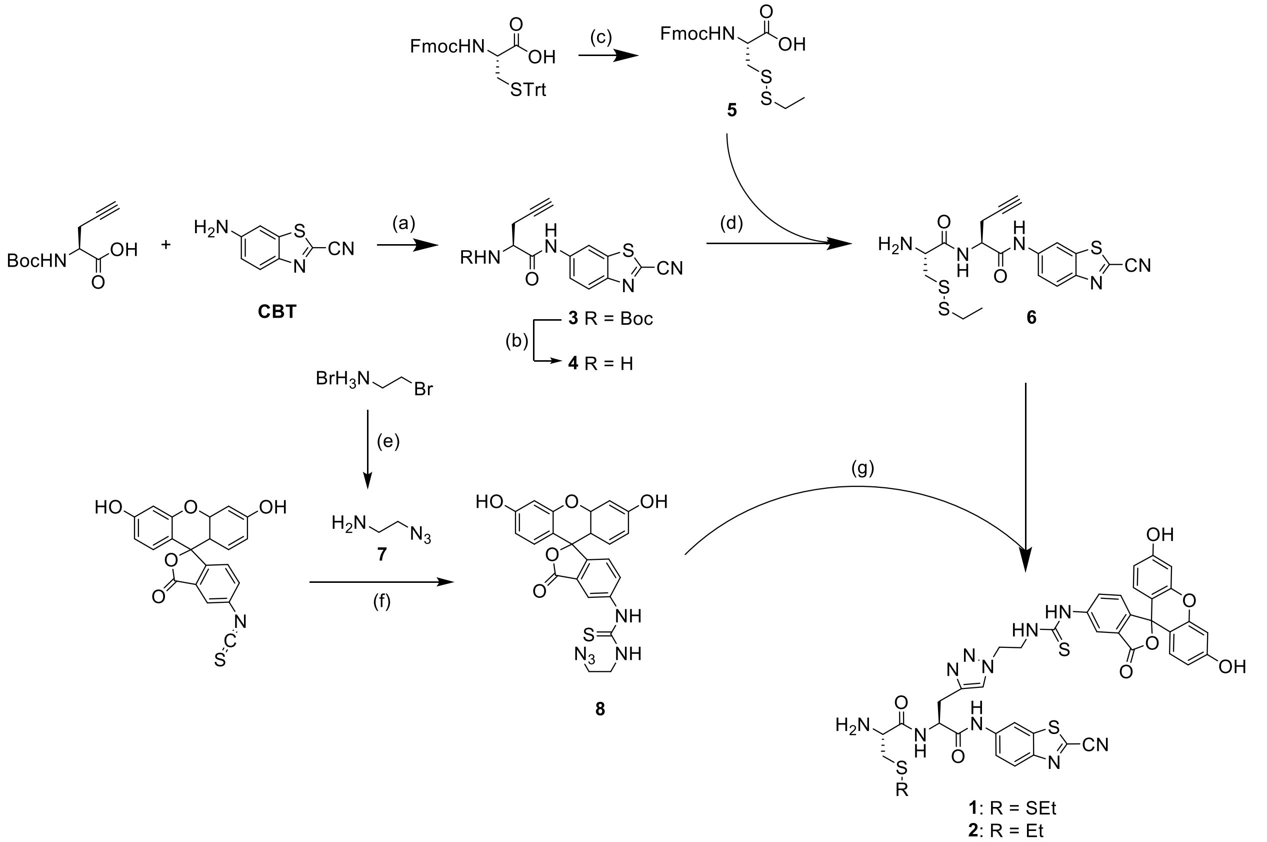

**Scheme S1.** Synthetic scheme of target compounds **1** and **2**. *Reagents and conditions:* (a) isobutyl chloroformate, N-methylmorpholine, THF, 0°C -> rt., OVN, 61%; (b) 20% TFA/DCM; (c) i) 10%TFA/DCM, 30 min; ii) PySSEt, MeOH, 30 min, 99% over two steps; (d) i) **5** or **Fmoc-Cys(Et)-OH**, HBTU, HOBt, DIPEA, DMF, r.t., 30 min; ii) 20% piperidine/DMF, rt., 5 min, HPLC, 64% over two steps; (e) $NaN_3$, $H_2O$, 80°C, 12h, 38%; (f) **7**, $NEt_3$, DMF, rt., 1.5h, 61%; (g) **6** or **6-ctrl**, $CuSO_4{\cdot}5H_2O$, $(BimC_4A)_3$, sodium ascorbate, DMF, rt. 3h, 50%.

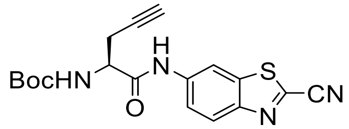

**tert-butyl (S)-(1-((2-cyanobenzo[d]thiazol-6-yl)amino)-1-oxopent-4-yn-2-yl)carbamate (3):** This compound was synthesized according to a literature procedure,[1] and the $^1$H NMR spectra was consistent with that reported. $^1$H NMR (300 MHz, Chloroform-*d*): δ/ppm = 8.62 (s, 1H), 8.06 (d, *J* = 8.9 Hz, 1H), 7.38 (d, *J* = 8.9 Hz, 1H), 5.42 (d, *J* = 7.6 Hz, 1H), 4.51 (d, *J* = 6.4 Hz, 1H), 2.81 (dddd, *J* = 42.4, 17.0, 6.2, 2.6 Hz, 2H), 2.15 (t, *J* = 2.6 Hz, 1H), 1.51 (s, 10H). $^{13}$C NMR (75 MHz, $CDCl_3$): δ/ppm = 169.16, 156.25, 148.69, 138.00, 136.72, 135.40, 125.26, 120.58, 112.94, 111.36, 103.36, 81.60, 78.71, 72.20, 53.64, 28.27, 21.54, 14.16. MS (ESI) calc'd for $C_{18}H_{18}N_4O_3S$: 393.0997 $[M+Na]^+$, found 393.1047.

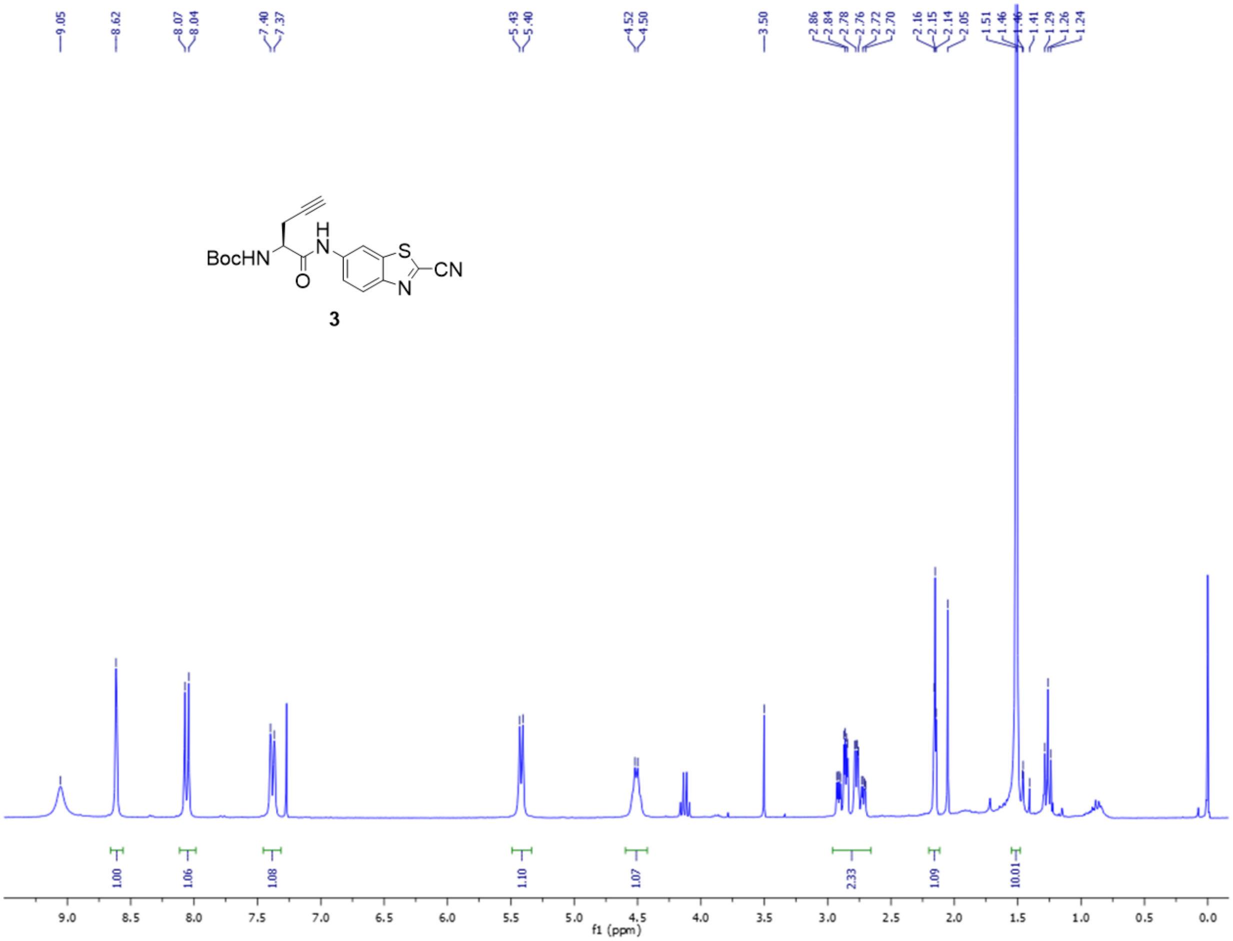

**Figure S6.** $^1$H-NMR spectrum of compound **3**.

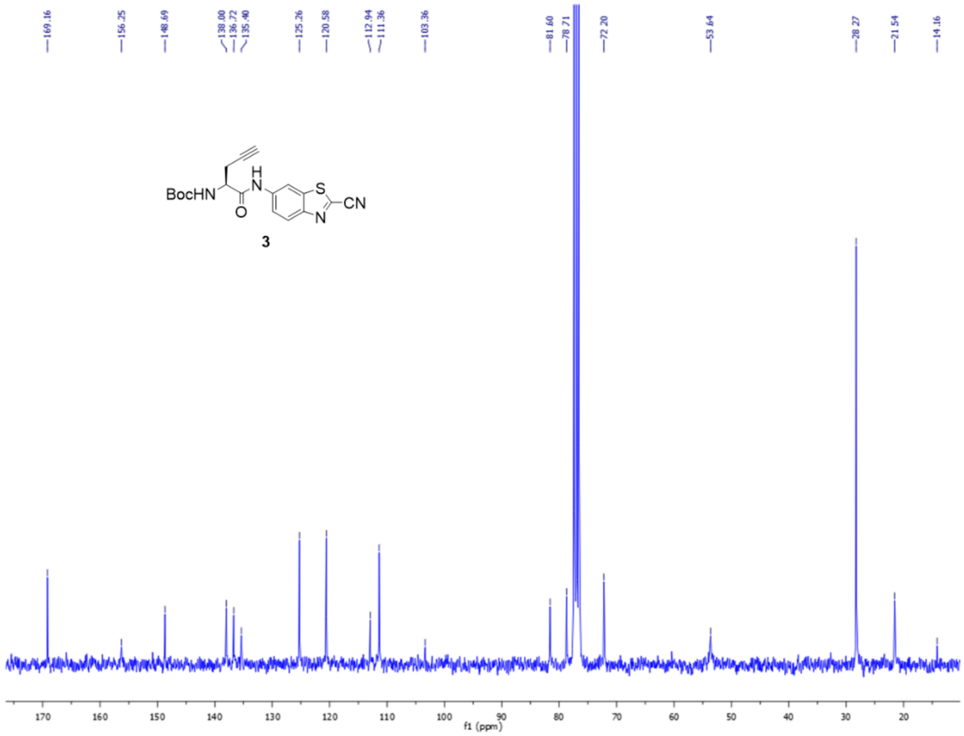

**Figure S7.** $^{13}$C-NMR spectrum of compound **3**.

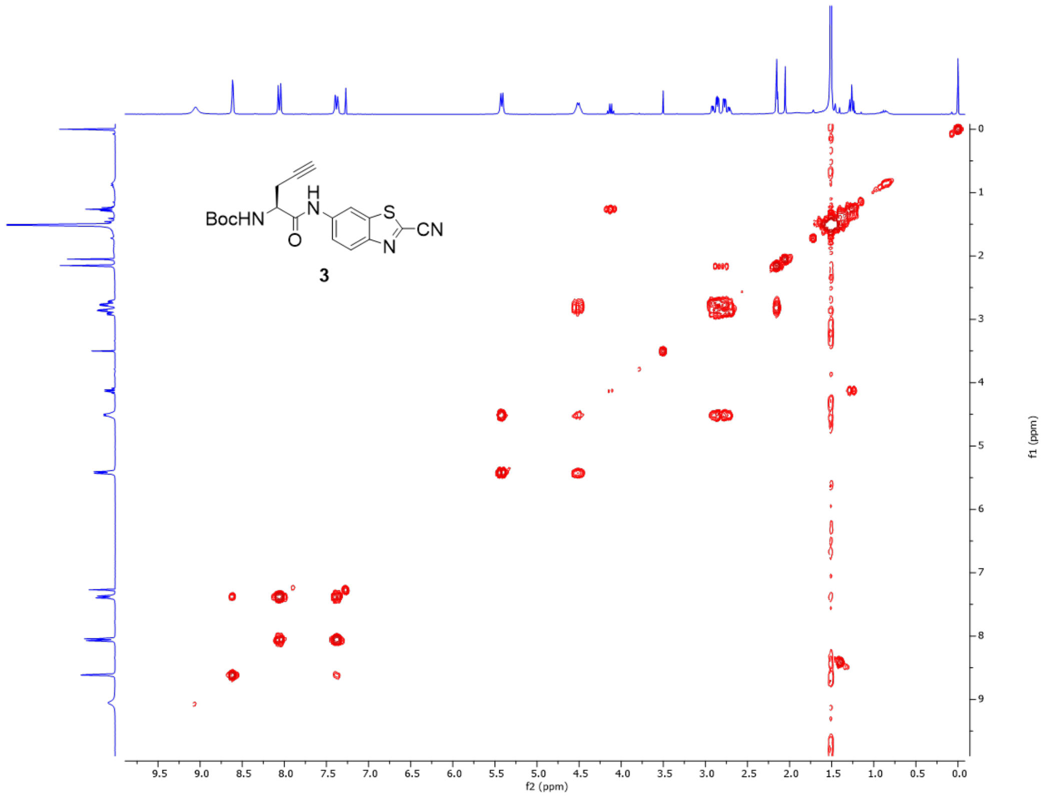

**Figure S8.** COSY spectrum of compound **3**.

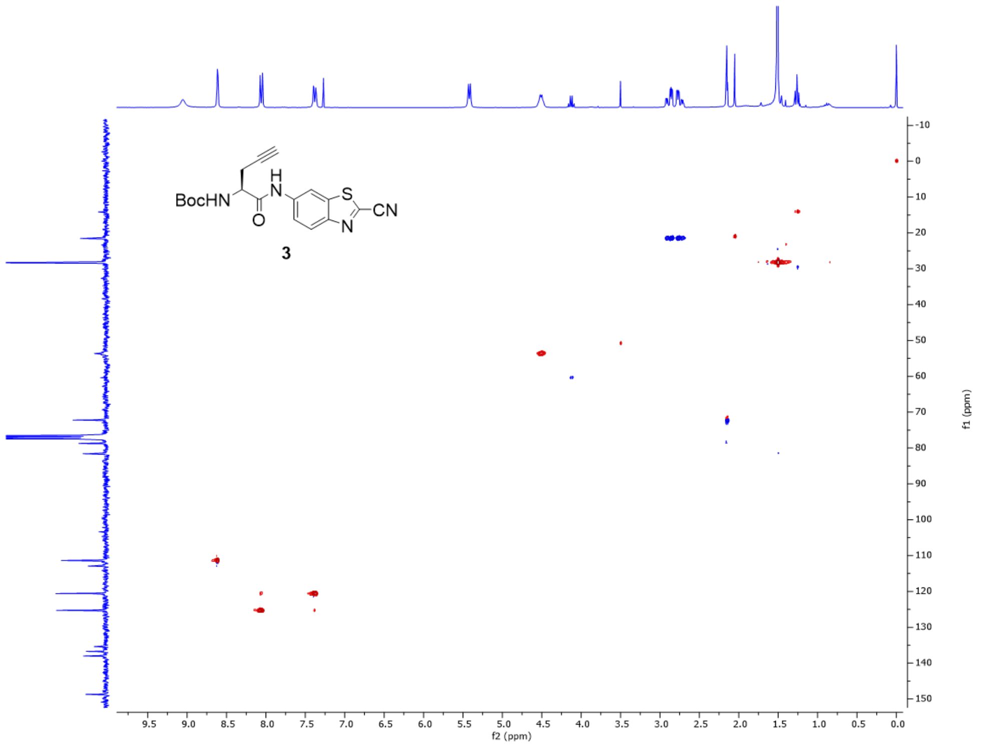

**Figure S8**. HSQC spectrum of compound **3**.

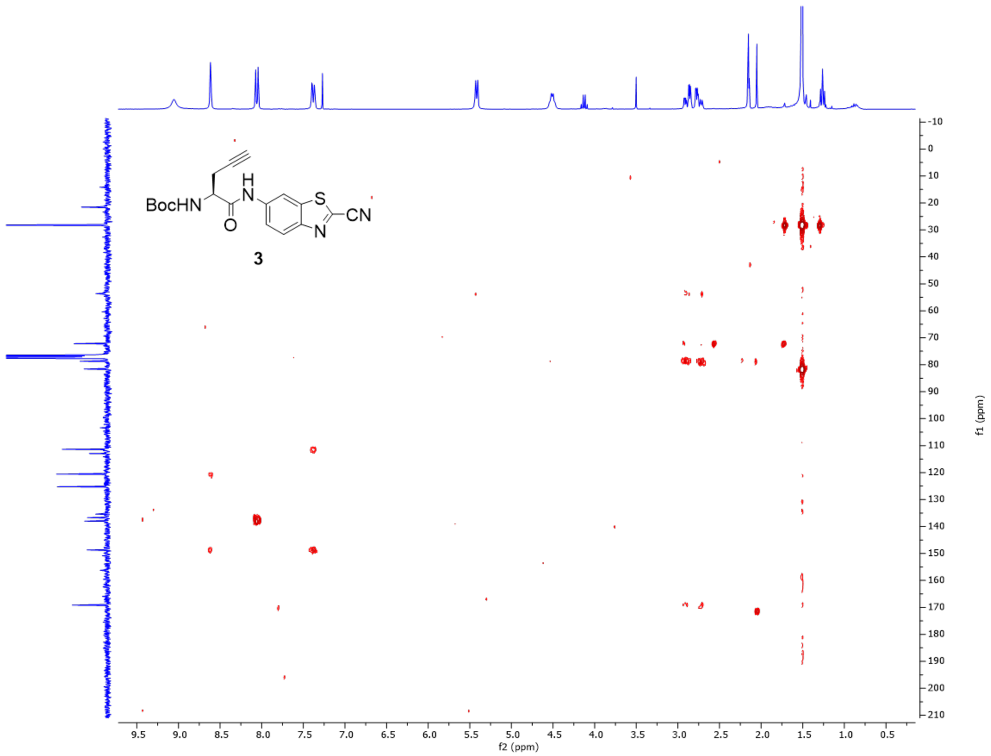

**Figure S9**. HMBC spectrum of compound **3**.

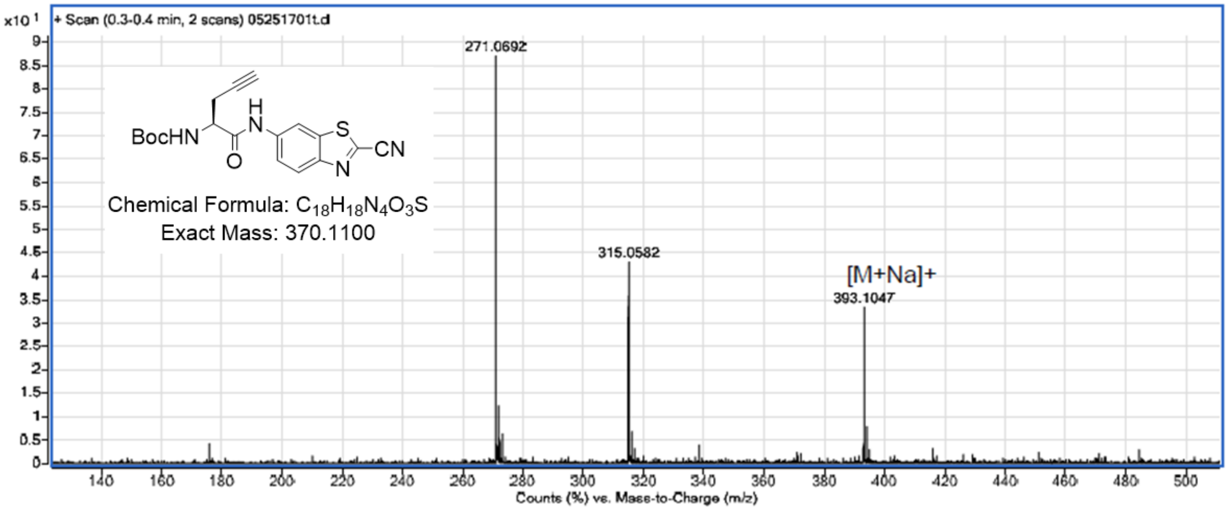

**Figure S10**. ESI-MS of compound **3**.

**(*S*)-2-amino-*N*-(2-cyanobenzo[d]thiazol-6-yl)pent-4-ynamide (4):** This compound was synthesized according to a literature procedure,[1] and the product was confirmed by comparing the $^1$H NMR spectra to that reported. $^1$H NMR (300 MHz, MeOD): δ/ppm = 8.70 (1 H, d, *J* 2.0), 8.16 (1 H, d, *J* 9.0), 7.74 (1 H, dd, *J* 9.0, 2.0), 4.26 (1 H, t, *J* 6.3), 2.98 (2 H, dd, *J* 6.2, 2.5), 2.70 (1 H, t, *J* 2.6). $^{13}$C NMR (75 MHz, MeOD) δ 167.35, 162.52, 162.04, 150.26, 139.77, 138.10, 137.33, 126.19, 122.14, 119.72, 113.99, 113.20, 77.00, 75.25, 53.67, 22.35. MS (ESI) calc'd for $C_{13}H_{10}N_4OS$: $[M+H]^+$ = 271.0645, found 271.0649.

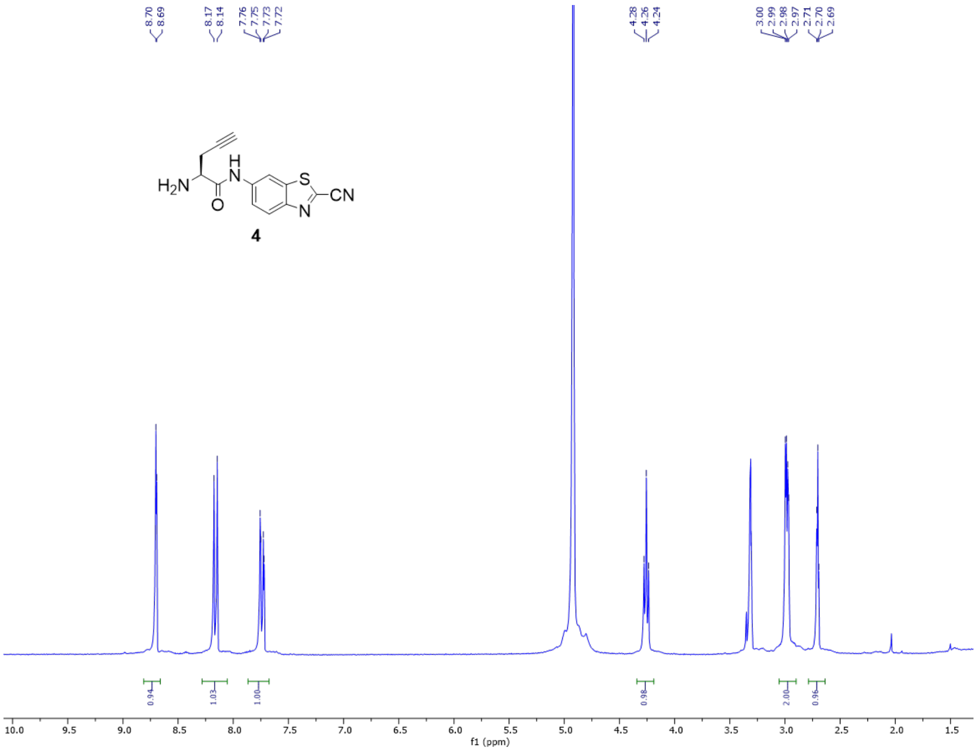

**Figure S11**. $^1$H-NMR spectrum of compound **4**.

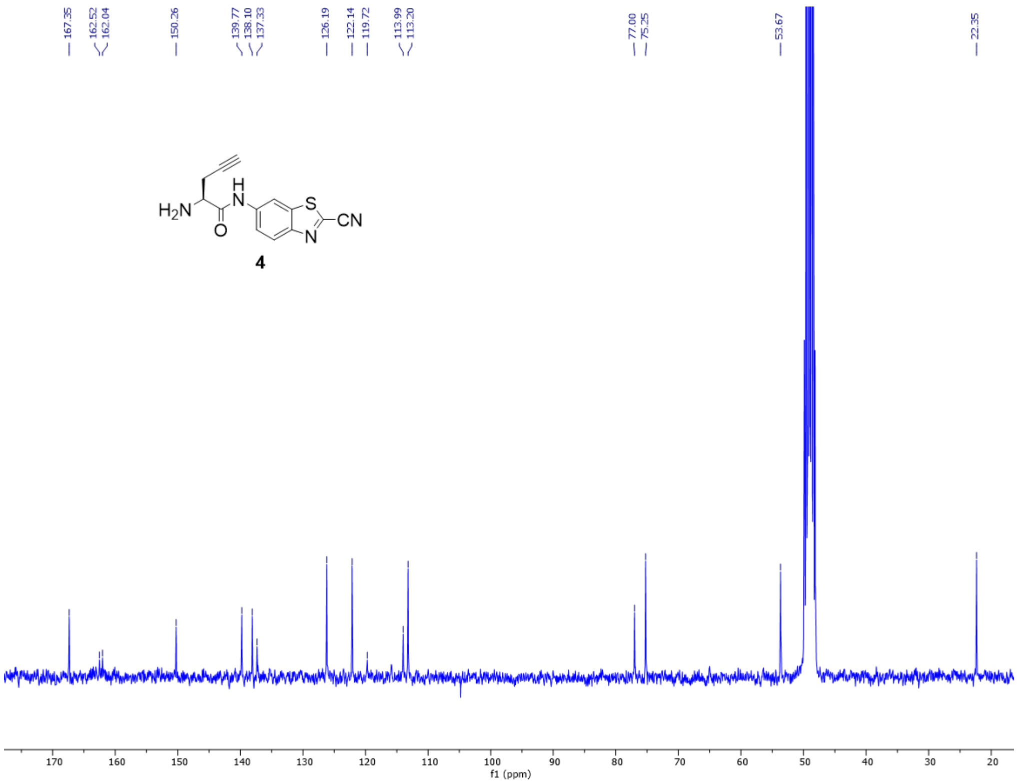

**Figure S12**. $^{13}$C-NMR spectrum of compound **4**.

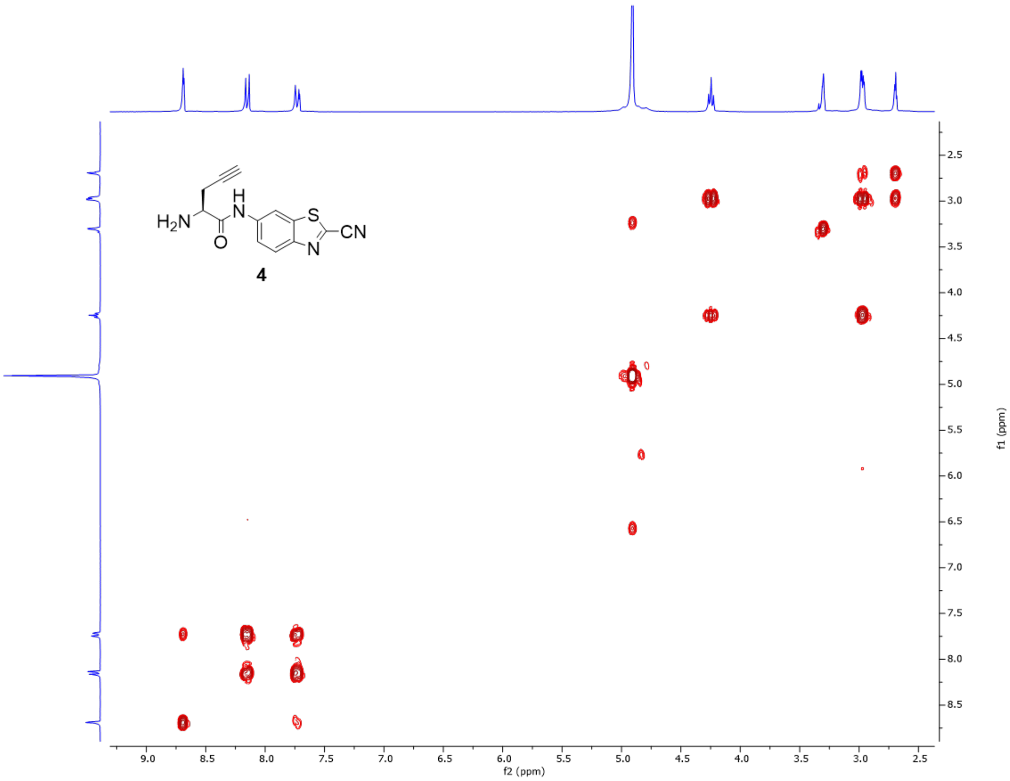

**Figure S13**. COSY spectrum of compound **4**.

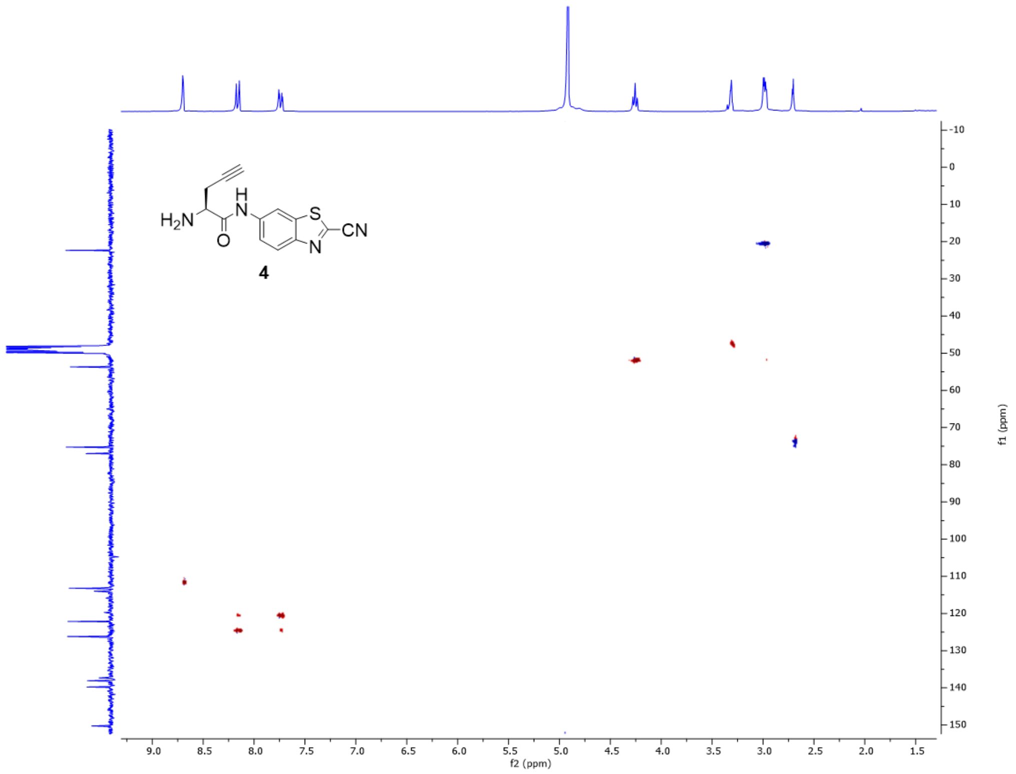

**Figure S14**. HSCQ spectrum of compound **4**.

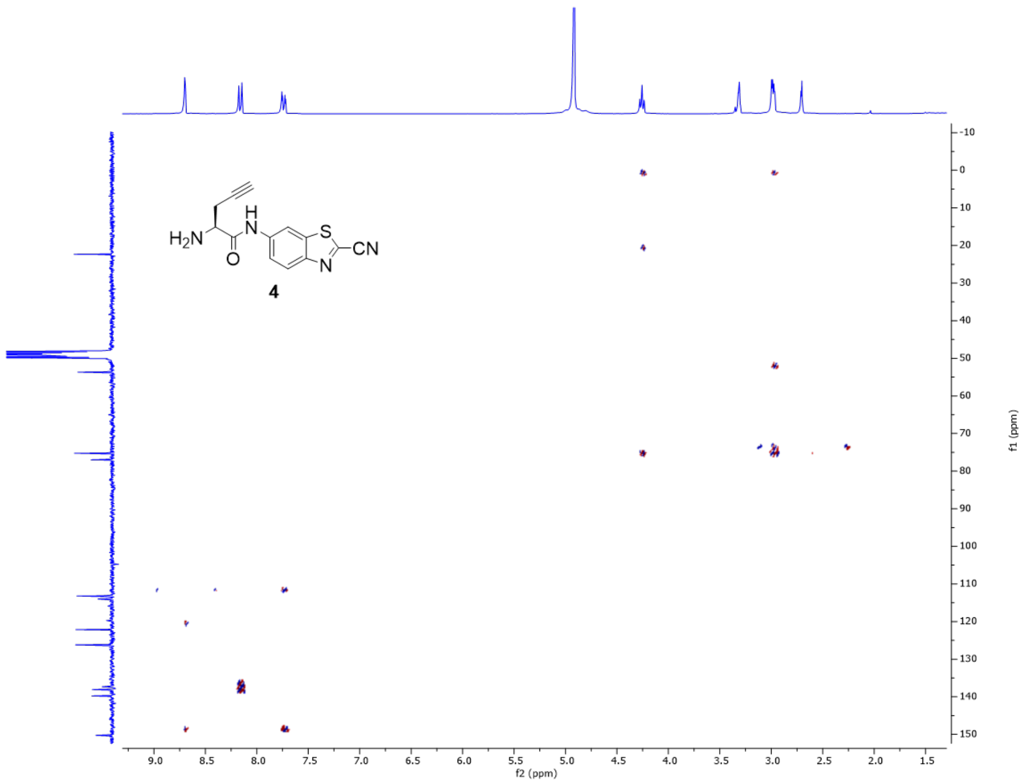

**Figure S15**. HMBC spectrum of compound **4**.

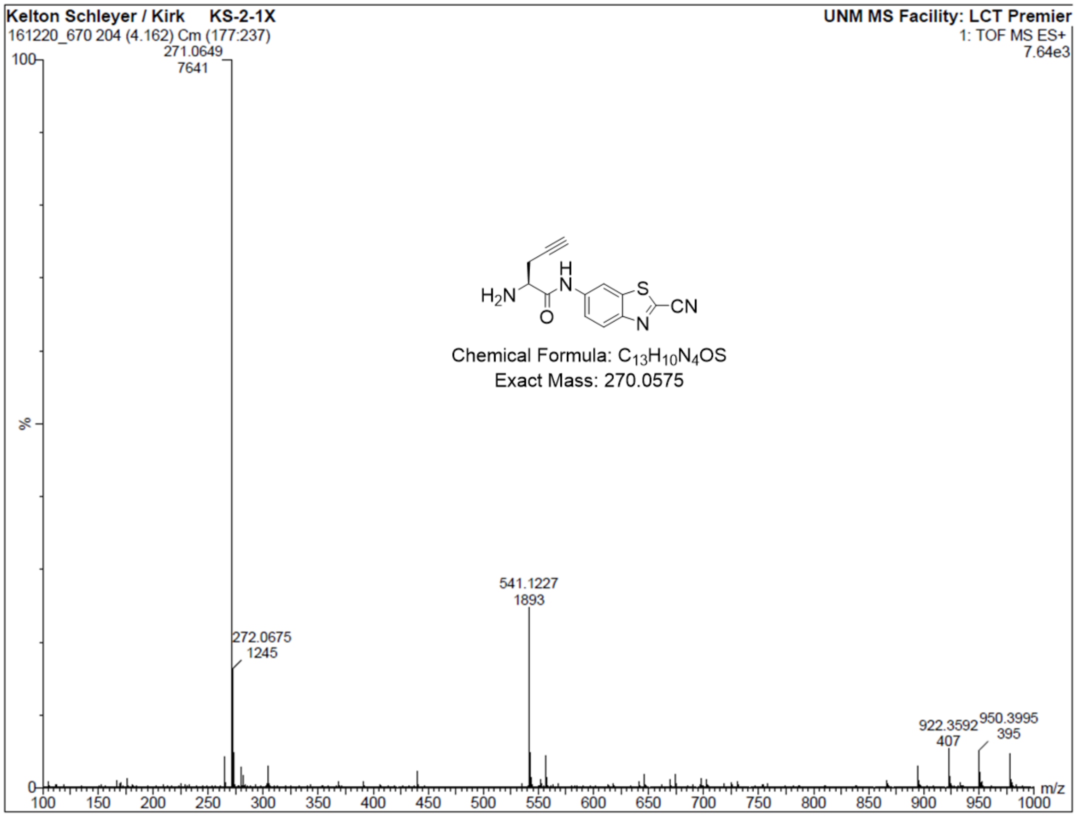

**Figure S16**. ESI-MS of compound **4**.

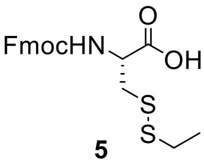

**5**

**N-(((9H-fluoren-9-yl)methoxy)carbonyl)-S-(ethylthio)-L-cysteine (5):**

Compound **5** was synthesized using a procedure adapted from the literature[1-2], and the $^{1}$H NMR spectra was compared to that of the literature references. To a solution of (Fmoc-Cys(Trt)-OH) (248 mg, 0.423 mmol) in dichloromethane (3.56 mL) was added triisopropylsilane (173 µL, 0.846 mmol, 2 eq.) and trifluoroacetic acid (0.440 mL, 10% v/v). Reaction was monitored by TLC ($R_f$ = 0.34 for product, hexanes/ethyl acetate = 1:2 with 1 drop HOAc). After 15min, solution was concentrated *in vacuo* to produce a clear viscous jelly.

The crude product was dissolved in methanol (5 mL) to give faint yellow solution, and PySSEt (145 mg, 0.846 mmol, 2 eq.) was added. The yellow color gradually intensified. Reaction was monitored by TLC ($R_f$ = 0.42 for product, hexanes/ethyl acetate = 1:2 with 1 drop HOAc). After 30 min reaction solution concentrated in vacuo, purified by flash chromatography (4:1 hexanes:ethyl acetate with 1.0% v/v acetic acid, isocratic); fractions combined, concentrated *in vacuo* to give viscous gel, which was diluted with methanol and toluene and concentrated again *in vacuo*. This process was repeated to co-evaporate acetic acid, and upon complete removal of AcOH and other solvents the pure product was afforded as a white powdered solid (153 mg, 90%). $^{1}$H-NMR (300 MHz, $CDCl_3$): δ/ppm = 7.76 (1 H, d, J 7.5), 7.63 (1 H, dd, J 16.1, 4.1), 7.40 (1 H, t, J 7.3), 7.32 (1 H, dd, J 10.7, 4.2), 5.70 (0 H, d, J 7.8), 4.74 (0 H, dd, J 12.7, 5.6), 4.42 (1 H, d, J 7.1), 4.24 (1 H, t, J 6.9), 3.21 (1 H, qd, J 14.2, 5.3), 2.71 (1 H, dd, J 14.6, 7.3), 1.38 – 1.18 (2 H, m). $^{13}$C-NMR (75 MHz, $CDCl_3$): δ/ppm = 175.14, 155.89, 143.61, 141.28, 127.75, 127.09, 125.10, 119.99, 67.41, 53.34, 47.03, 40.25, 32.62, 14.28.

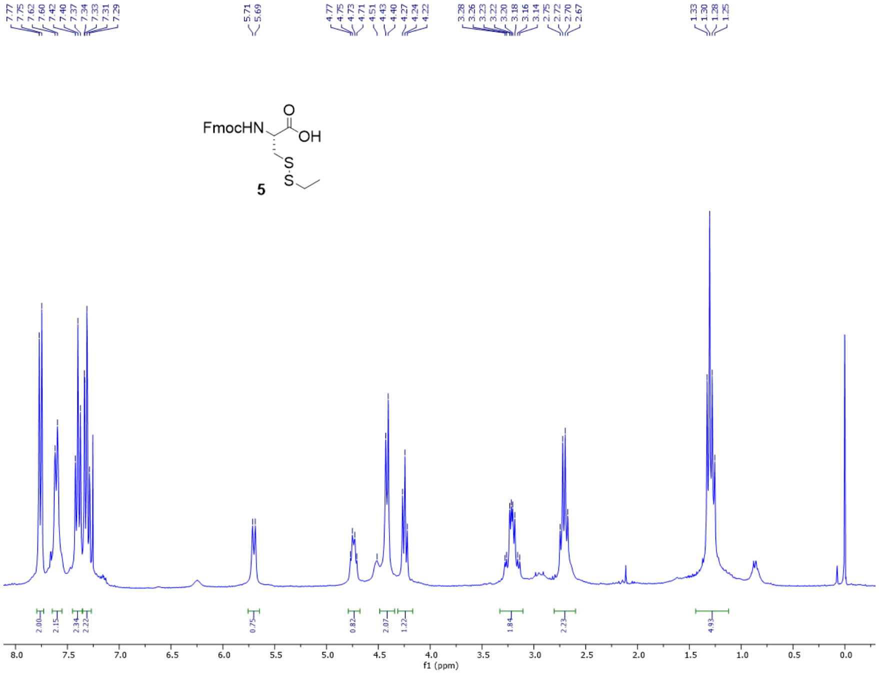

**Figure S17**. $^{1}$H-NMR spectrum of compound **5**.

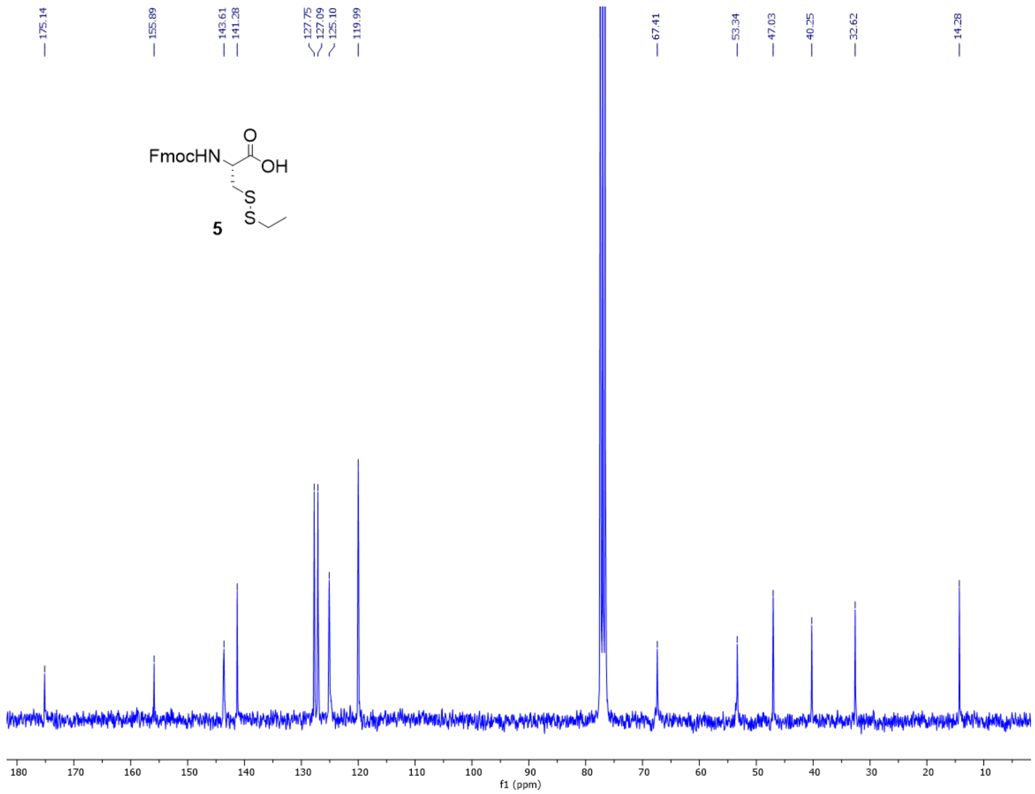

**Figure S18**. $^{13}$C-NMR spectrum of compound **5**.

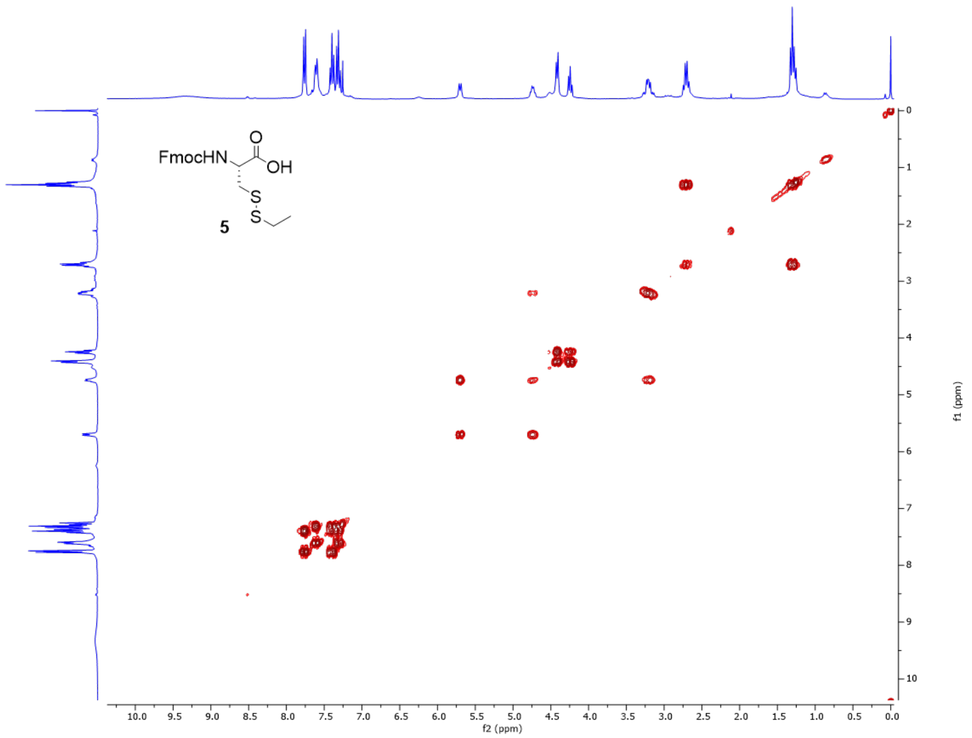

**Figure S19**. COSY spectrum of compound **5**.

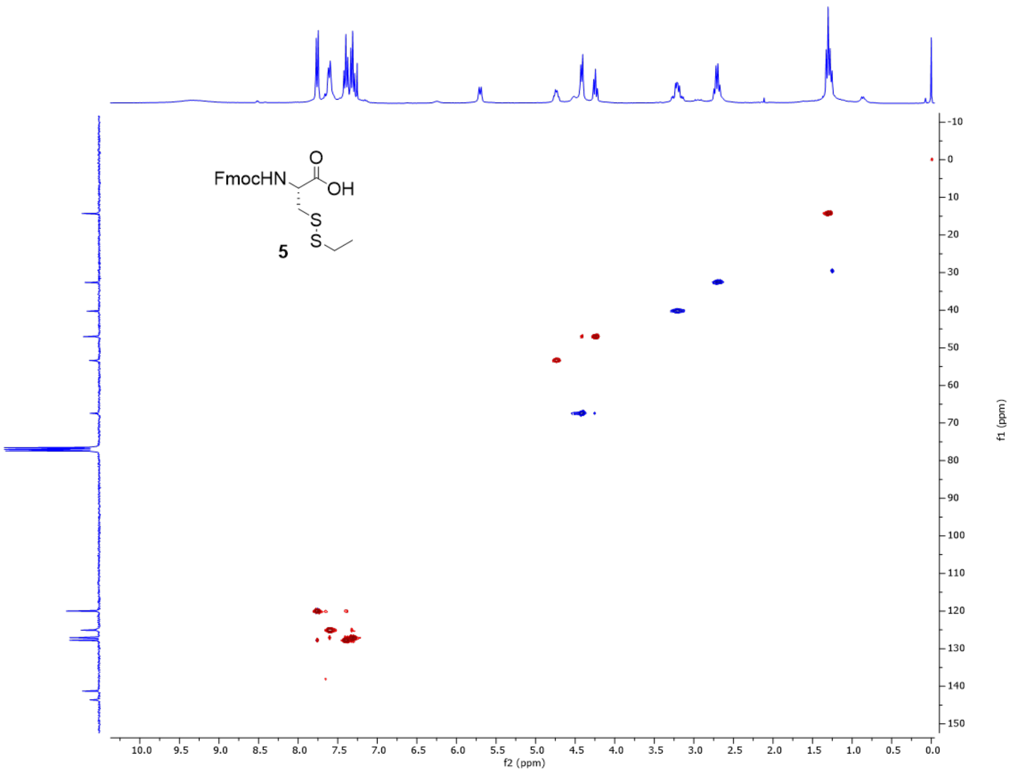

**Figure S20**. HSQC spectrum of compound **5**.

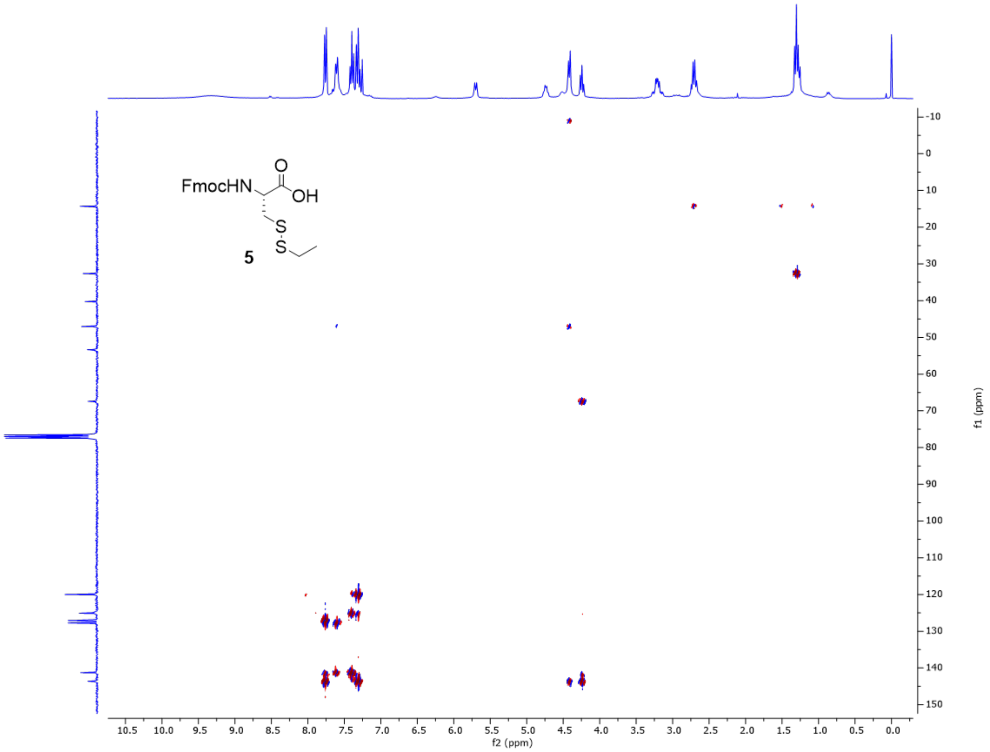

**Figure S21**. HMBC spectrum of compound **5**.

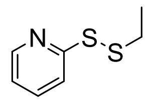

**2-(ethyldisulfaneyl)pyridine:** This reagent was synthesized according to a literature procedure[3] and the identity was confirmed by comparison of the $^{1}H$ NMR spectrum.

**(S)-2-((R)-2-amino-3-(ethyldisulfanyl)propanamido)-N-(2-cyanobenzo[d]thiazol-6-yl)pent-4-ynamide (6):**

To a solution of **3** (18 mg, 0.0437 mmol, 1 eq.) in DMF in a r.b. flask was added HOBt (20 mg, 0.1311 mmol, 3 eq.), then HBTU (50 mg, 0.1311 mmol, 3 eq.). The solution was stirred for 30 min, after which was added **2** (13 mg, 0.048 mmol, 1.1 eq.). The solution was stirred an additional 30 min, after which DIPEA (23 uL, 0.131 mmol, 3 eq.) was added. The clear solution gradually turned translucent yellow in color. Reaction progress was monitored by HPLC ($t_R$ = 12.88 min for product). Upon complete conversion (~30 min) the reaction solution was transferred to a separatory funnel, diluted with EA (50 mL) and washed with DI water (6 x 20 mL) until HPLC confirmed coupling byproducts were removed from the organic layer. The organic layer was washed with brine (3 x 20 mL) then dried over sodium sulfate. Crude product concentrated under vacuum to produce a yellow solid, used directly in the next step.

The crude product was re-dissolved in DMF (2.4 mL) to give yellow solution, then piperidine (600 uL, giving a 20% solution) was added. The solution immediately turned deeper yellow in color. After 5 min the solution was transferred to a separatory funnel, diluted with ethyl acetate, and washed with water (3 x 40 mL) then brine (3 x 30 mL). The organic layer was dried over sodium sulfate, concentrated in vacuo to give a yellow and white solid. The crude product was dissolved in 50% MeOH/$H_2O$ to give a yellow solution with prominent white suspended particles (white compound suspected to be dibenzofulvene-piperidine adduct, poor water solubility). The suspension was centrifuged and the supernatant was purified by HPLC to afford the pure product (11 mg, 58% over two steps). $^{1}H$-NMR (500 MHz, MeOD): δ/ppm = 8.67 (1 H, d, J 1.9), 8.14 (1 H, d, J 9.0), 7.72 (1 H, dd, J 9.0, 2.0), 4.76 (1 H, t, J 7.0), 4.27 (1 H, dd, J 8.6, 4.6), 3.35 (2 H, dt, J 9.5, 4.7), 3.03 (1 H, dd, J 14.6, 8.6), 2.87 – 2.67 (6 H, m), 2.48 (1 H, t, J 2.5), 1.30 (4 H, dd, J 9.2, 5.4). $^{13}C$-NMR (126 MHz, MeOD): δ/ppm = 170.41, 168.71, 150.03, 140.31, 138.09, 136.98, 129.70, 128.84, 126.06, 122.21, 114.04, 112.94, 79.62, 72.95, 54.79, 53.08, 39.55, 32.79, 22.83, 14.42. MS (ESI) calc'd for $C_{18}H_{19}N_5O_2S_3$: 434.0771 $[M+H]^+$, found 434.0800; 456.0598 $[M+Na]^+$, found 456.0628.

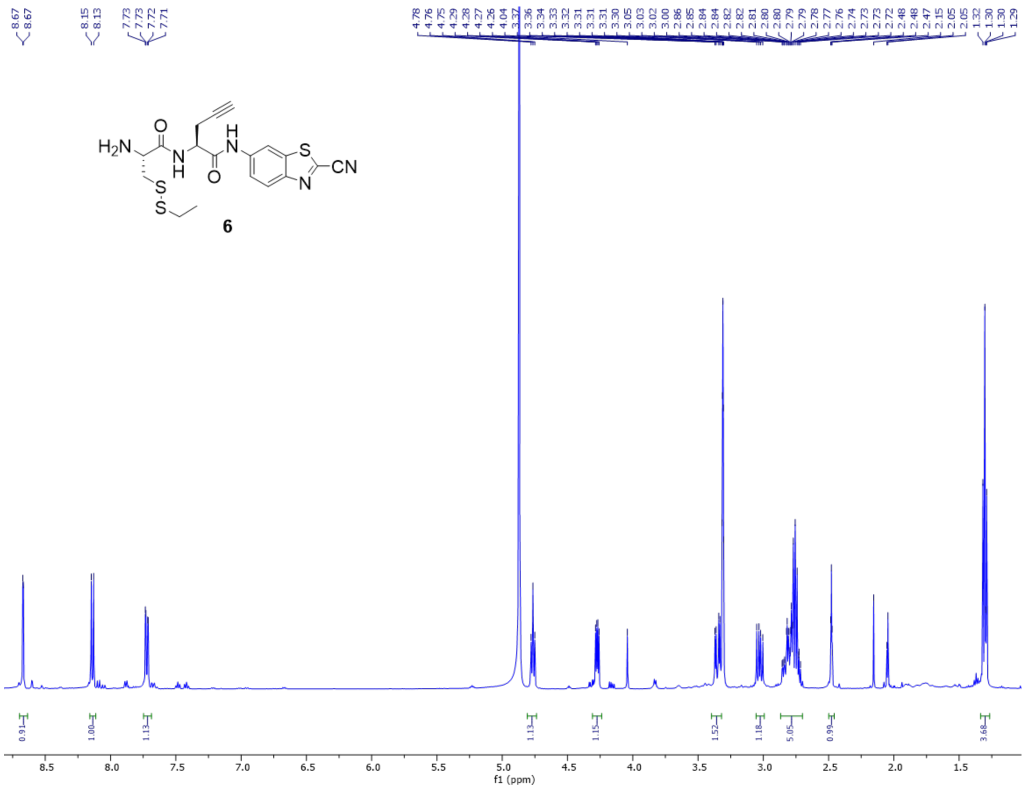

**Figure S22**. [1]H-NMR spectrum of compound **6**.

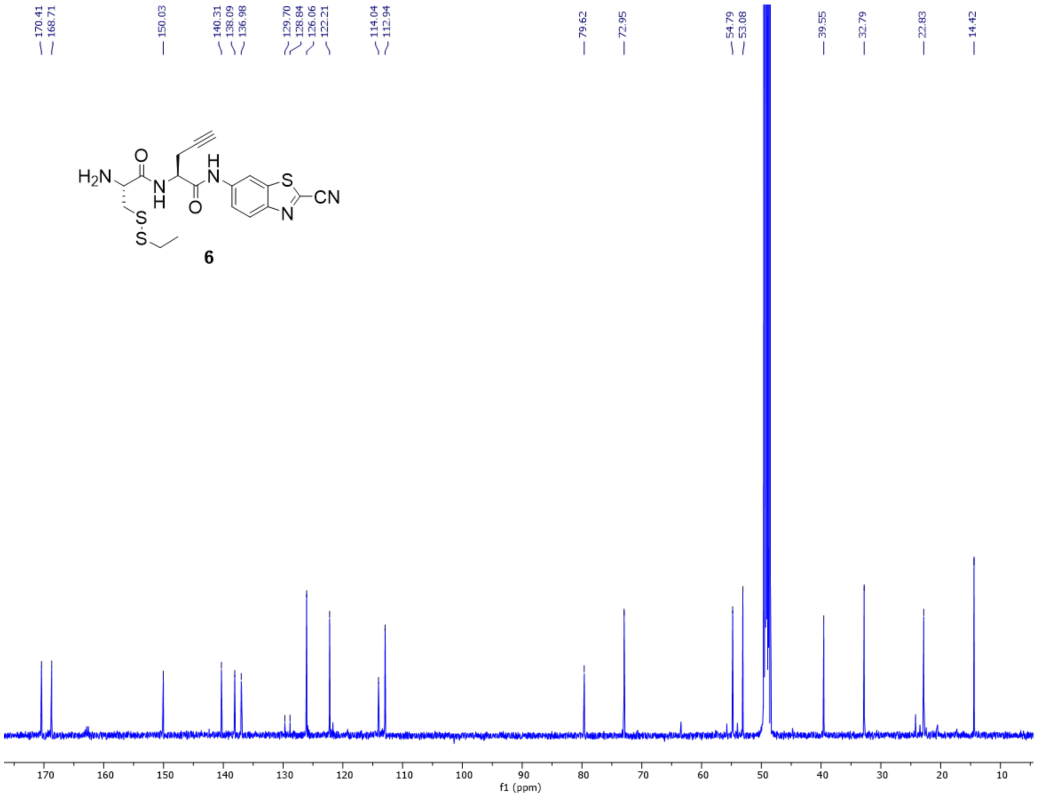

**Figure S23**. [13]C-NMR spectrum of compound 6.

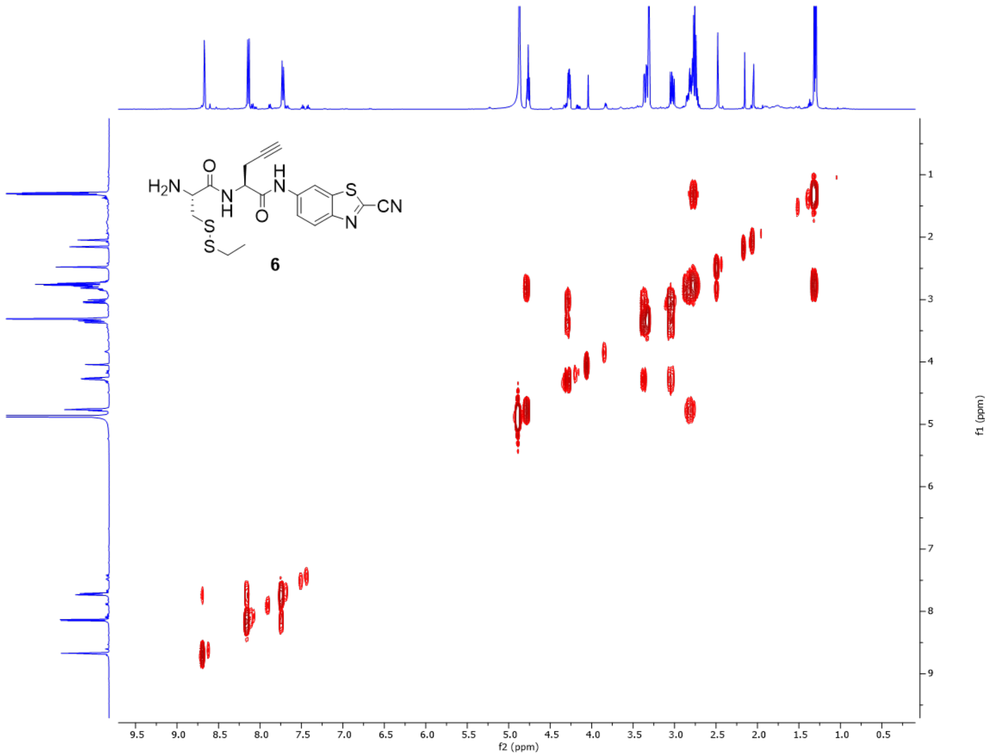

**Figure S24**. COSY spectrum of compound **6**.

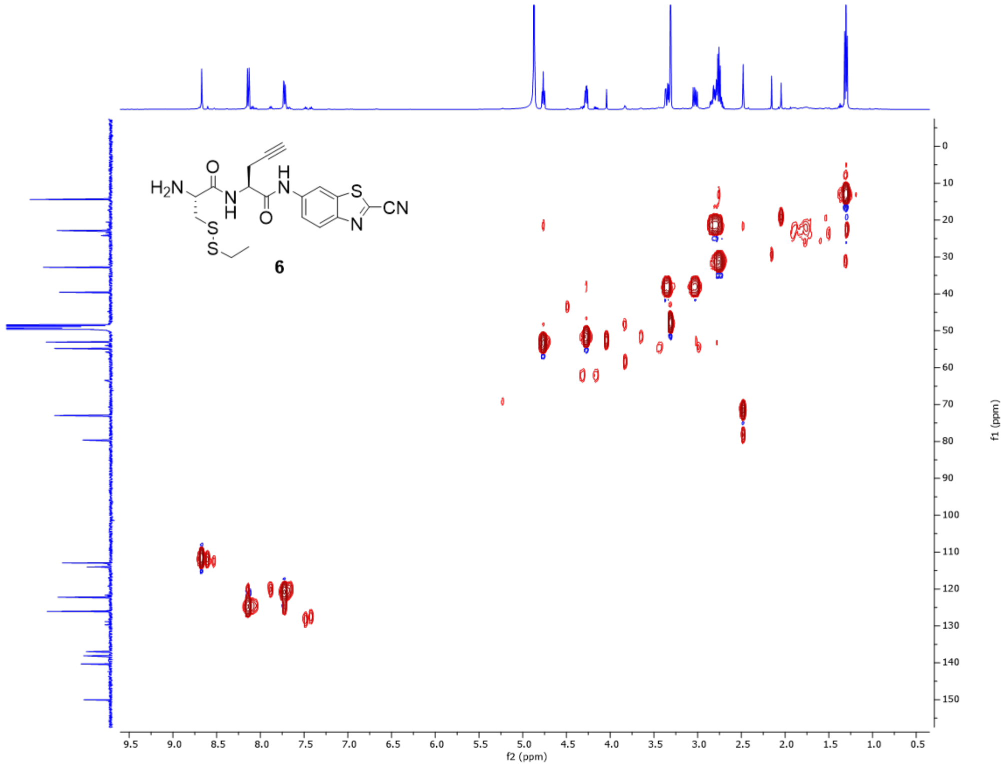

**Figure S25**. HSQC spectrum of compound **6**.

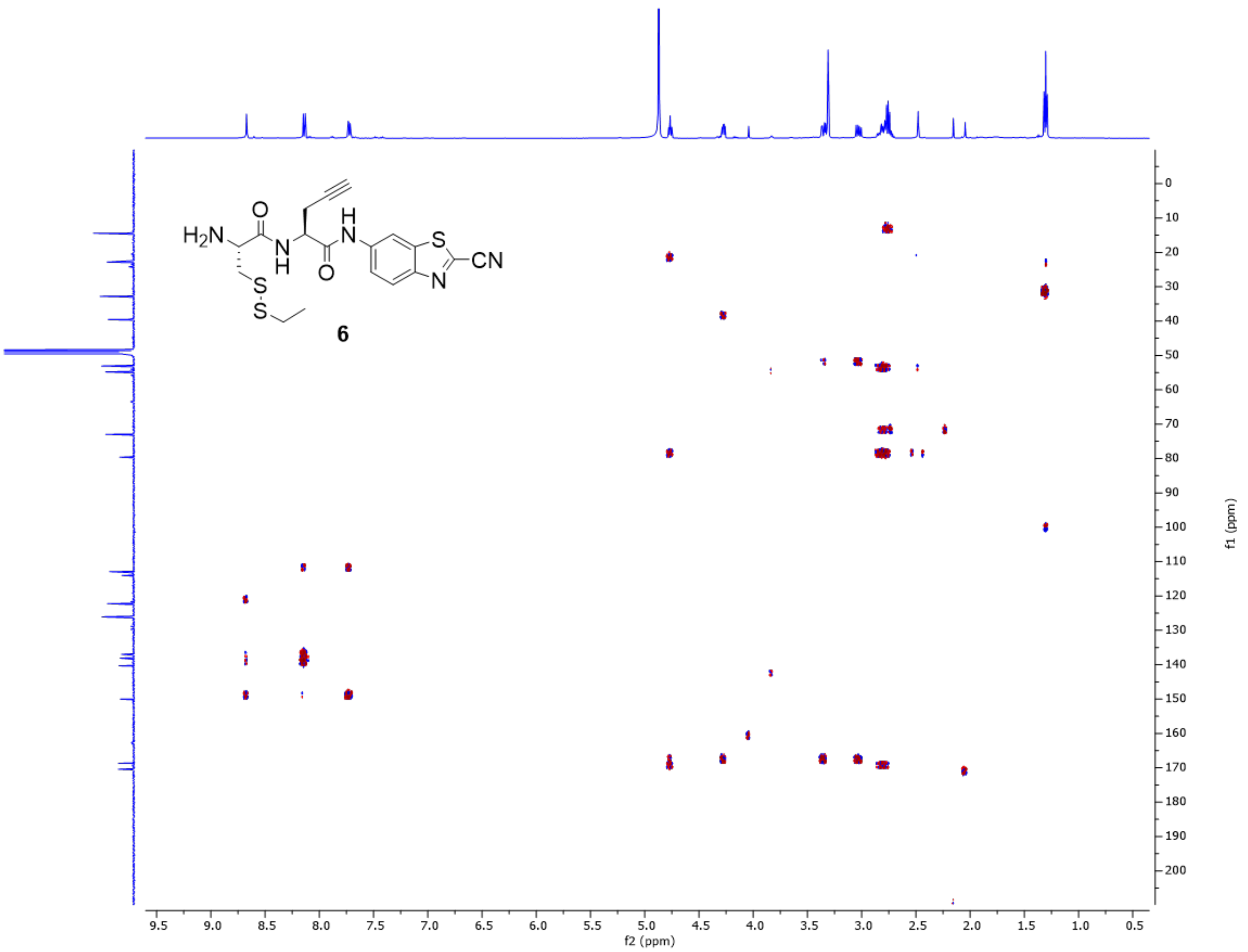

**Figure S26**. HMBC spectrum of compound **6**.

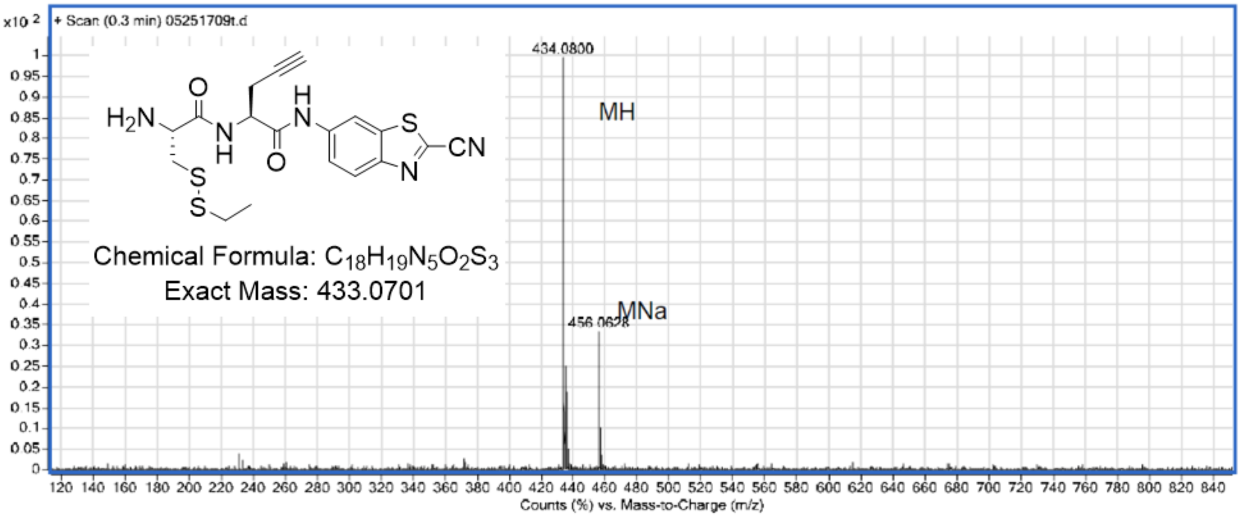

**Figure S27**. ESI-MS of compound **6**.

**(S)-2-((R)-2-amino-3-(ethylthio)propanamido)-N-(2-cyanobenzo[d]thiazol-6-yl)pent-4-ynamide (6-ctrl):** Synthesized using the same procedure as that for the synthesis of **4**. For synthesis of **4-ctrl**, **Fmoc-Cys(Et)-OH** was used instead of compound **3**; all other methods were identical. $^{1}H$ NMR (600 MHz, Methanol-$d_4$): δ/ppm = 8.67 (d, $J$ = 2.1 Hz, 1H), 8.15 (d, $J$ = 9.0 Hz, 1H), 7.72 (dd, $J$ = 9.0, 2.1 Hz, 1H), 4.76 (t, $J$ = 7.0 Hz, 1H), 4.09 (dd, $J$ = 8.6, 4.8 Hz, 1H), 3.18 (dd, $J$ = 14.6, 4.8 Hz, 1H), 2.93 – 2.87 (m, 1H), 2.86 – 2.73 (m, 2H), 2.63 (q, $J$ = 7.4 Hz, 2H), 2.48 (t, $J$ = 2.7 Hz, 1H), 1.24 (t, $J$ = 7.4 Hz, 3H). MS (ESI) calc'd for $C_{18}H_{19}N_5O_2S_2$:[M+H]$^{+}$ = 402.1050, found 402.1063; [M+Na]$^{+}$ = 424.0877, found 424.0932.

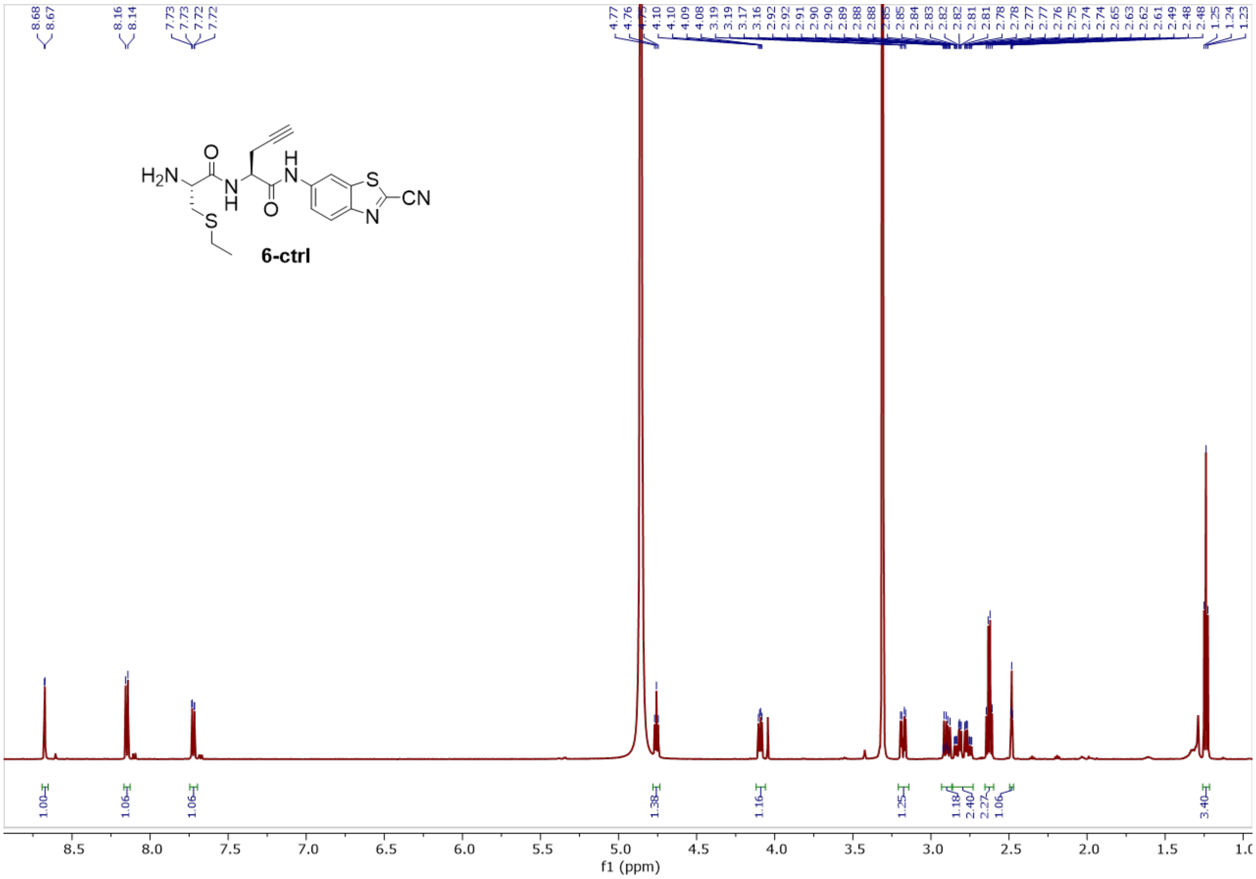

**Figure S28**. $^{1}H$-NMR spectrum of compound **6-ctrl**.

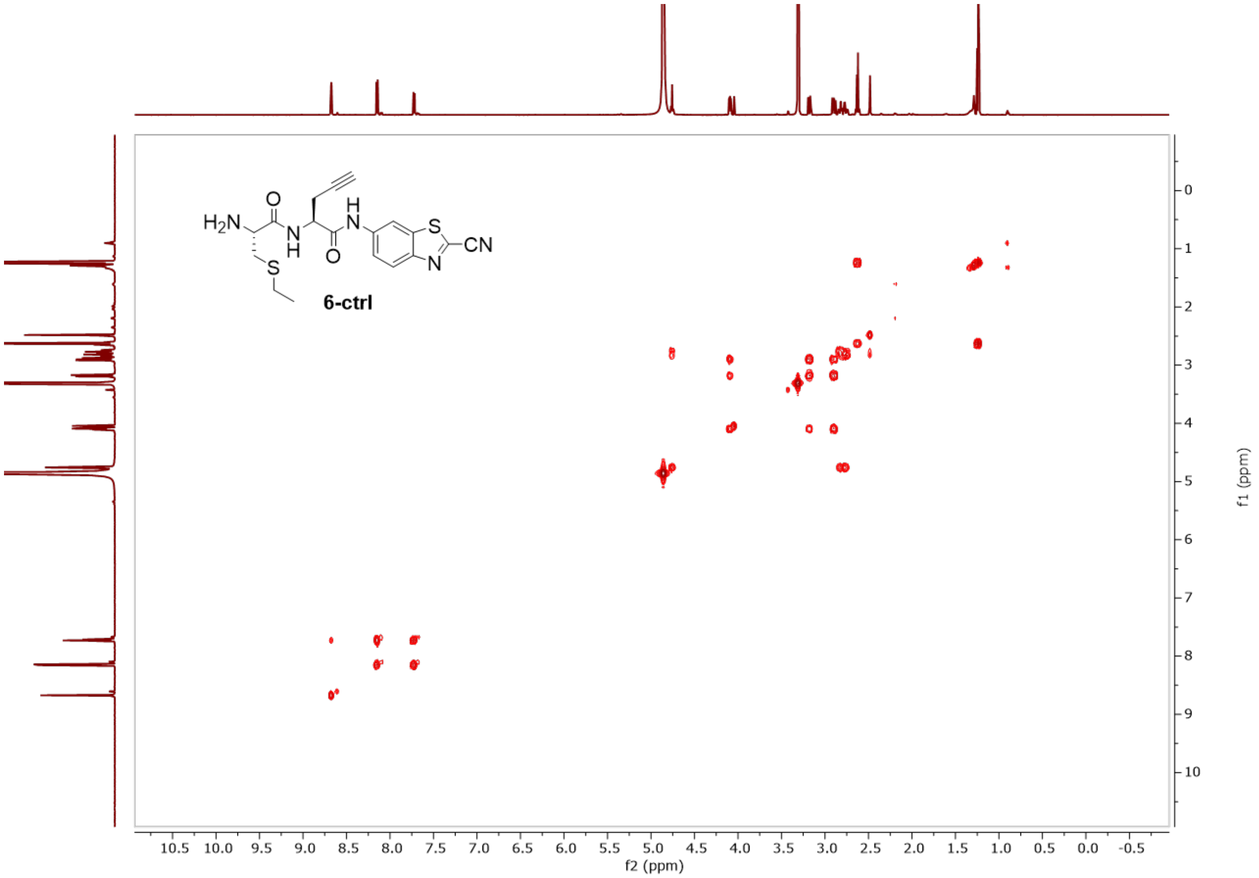

**Figure S29**. COSY spectrum of compound **6-ctrl**.

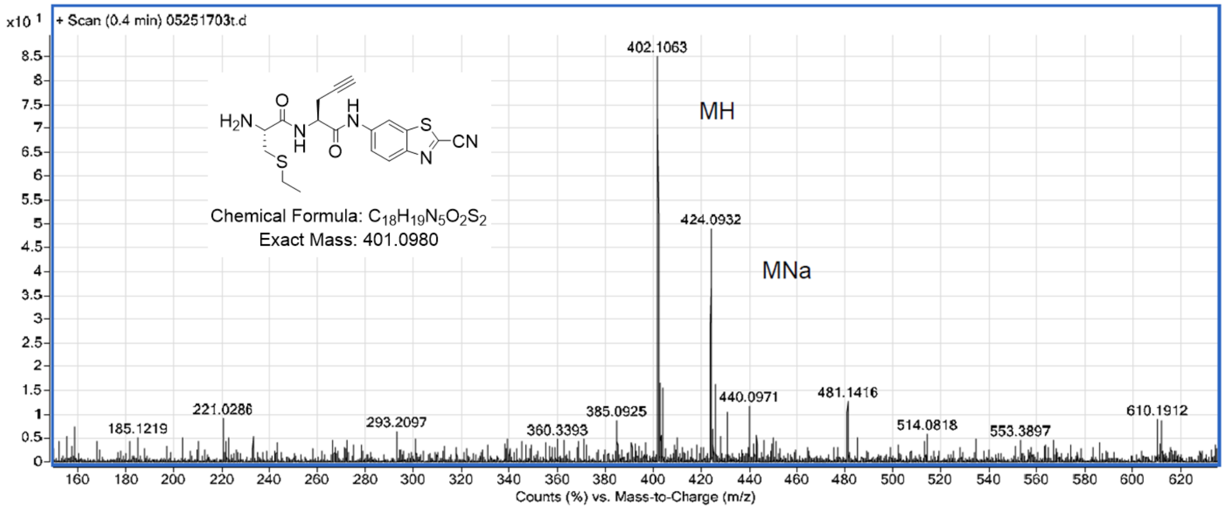

**Figure S30**. ESI-MS spectrum of compound **6-ctrl**.

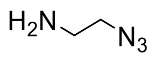

**2-azidoethylamine (7):** This compound was synthesized according to a literature procedure,[4] and the $^1$H NMR spectrum was consistent with that reported.

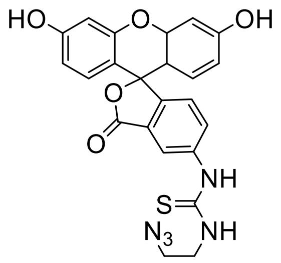

**1-(2-azidoethyl)-3-(3',6'-dihydroxy-3-oxo-3H-spiro[isobenzofuran-1,9'-xanthen]-5-yl)thiourea (8):** Adapted from reference[5-6]. FITC-I (30 mg, 0.077 mmol, 1 eq.) dissolved in dry DMF (3 mL total) to give a yellow solution. 2-azidoethylamine (10 mg, 0.1155 mmol, 1.5 eq.) in dry DMF was added to the reaction solution, which immediately turned orange. Triethylamine (22 uL, 0.154 mmol, 2.0 eq.) was added and the reaction was covered in tin foil. Stirred for 2.5h, after which HPLC (sample diluted in DMF) indicated complete consumption of the SM ($t_R$ = 11.97 min) and the presence of one new peak ($t_R$ = 11.71 min). This new peak was missing a shoulder lambda-max = 276 nm, which is present in FITC-I. The reaction mixture was centrifuged and the compound was purified by HPLC to give an orange solid, 13.7 mg (37%). $^1$H-NMR (500 MHz, MeOD): δ/ppm = 8.32 (1 H, d, J 1.5), 8.27 (3 H, d, J 1.2), 7.96 (1 H, dd, J 8.2, 1.9), 7.87 (3 H, dd, J 8.2, 1.8), 7.29 (1 H, d, J 8.2), 7.24 (3 H, d, J 8.3), 7.05 – 6.92 (8 H, m), 6.88 (7 H, d, J 8.6), 6.80 – 6.69 (8 H, m), 3.83 (5 H, t, J 5.6), 3.61 (5 H, t, J 5.8). $^{13}$C-NMR (126 MHz, MeOD): δ/ppm = 183.45, 183.45, 142.73, 142.73, 131.62, 131.62, 116.02, 116.02, 103.44, 103.44, 51.18, 44.71. MS (ESI) calc'd for $C_{23}H_{17}N_5O_5S$: 476.1020 $[M+H]^+$, found 476.1036; 474.0880 $[M-H]^-$, found 474.0875.

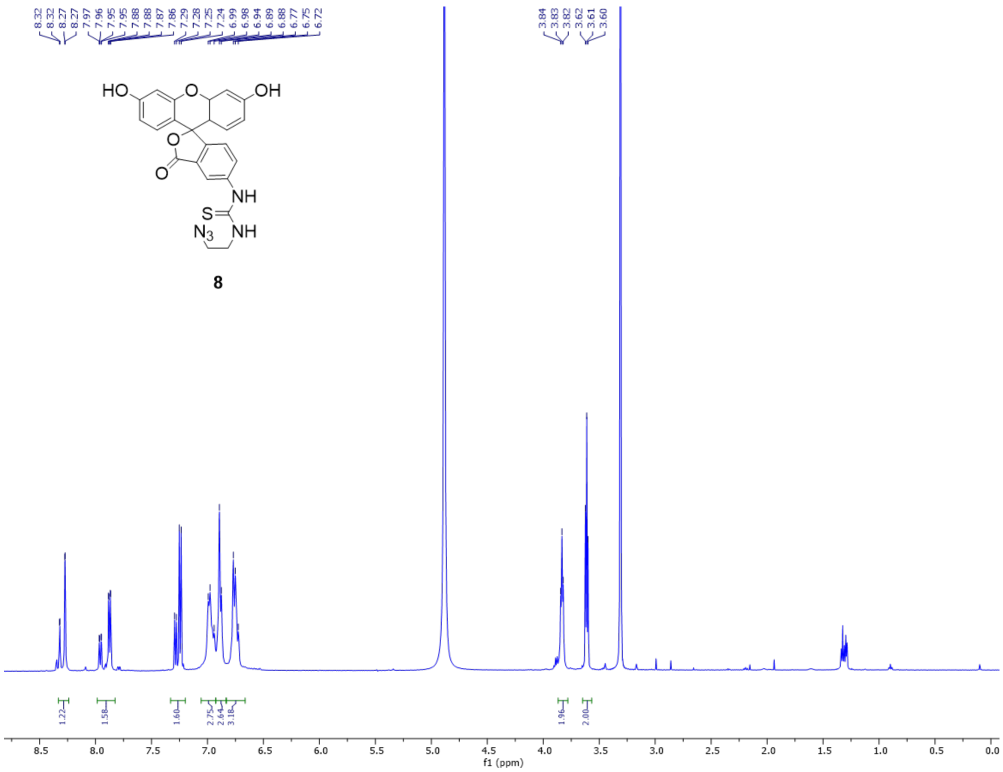

**Figure S31**. $^{1}$H-NMR spectrum of compound **8**.

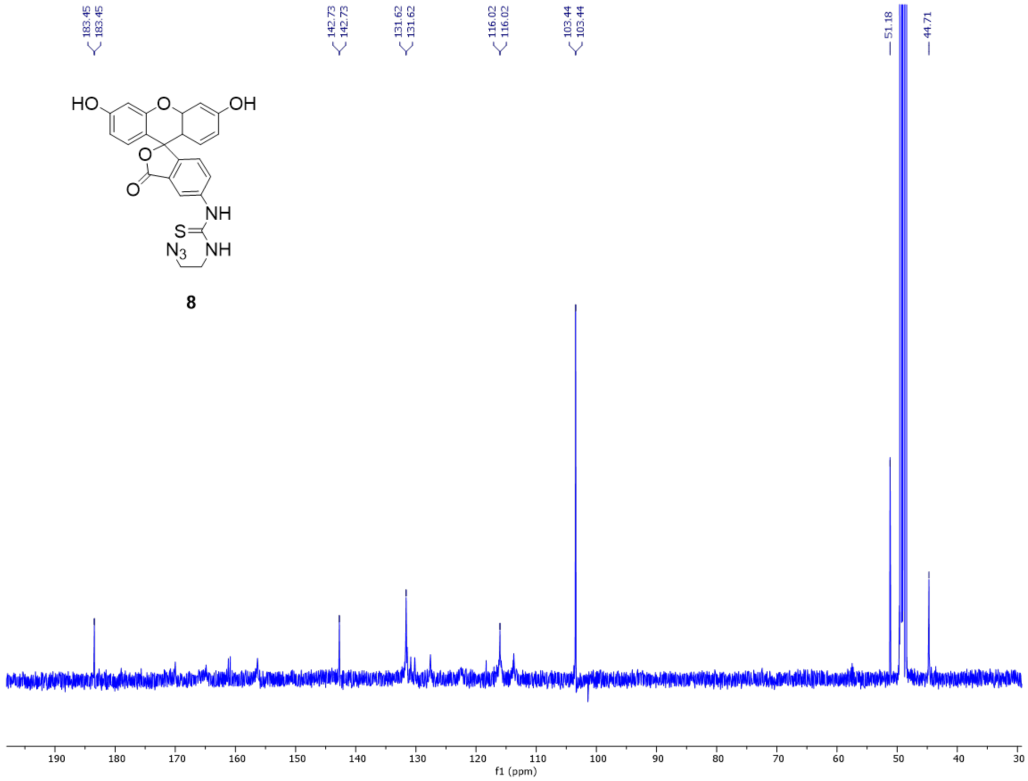

**Figure S32**. $^{13}$C-NMR spectrum of compound **8**.

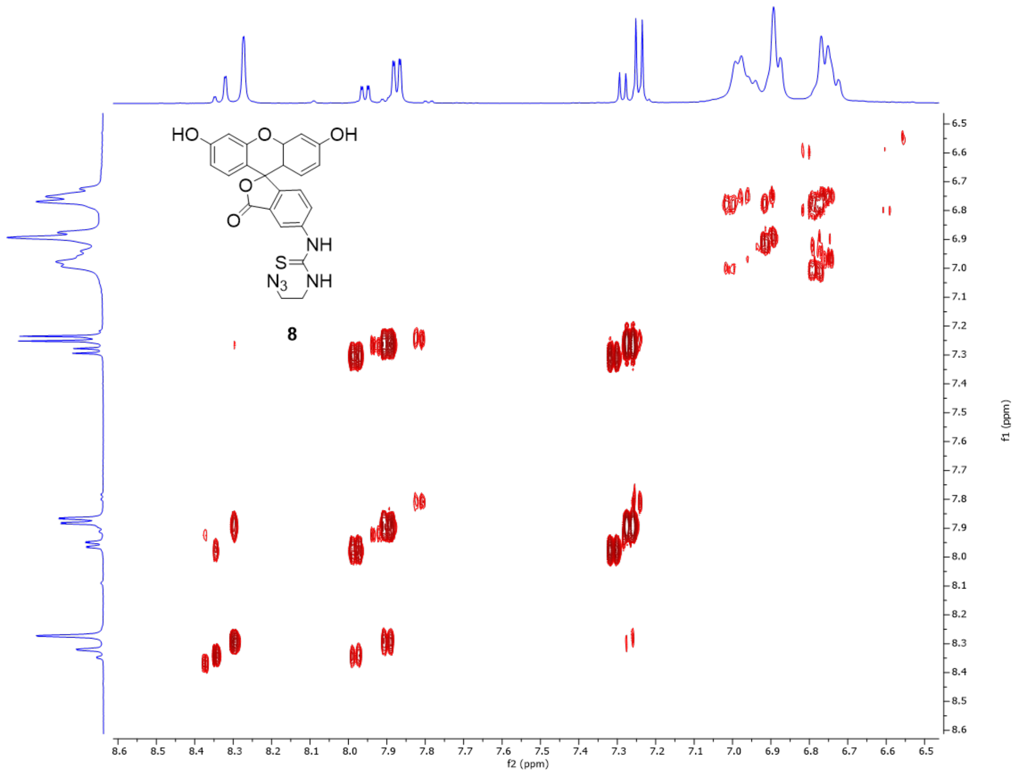

**Figure S33**. COSY spectrum of compound **8**.

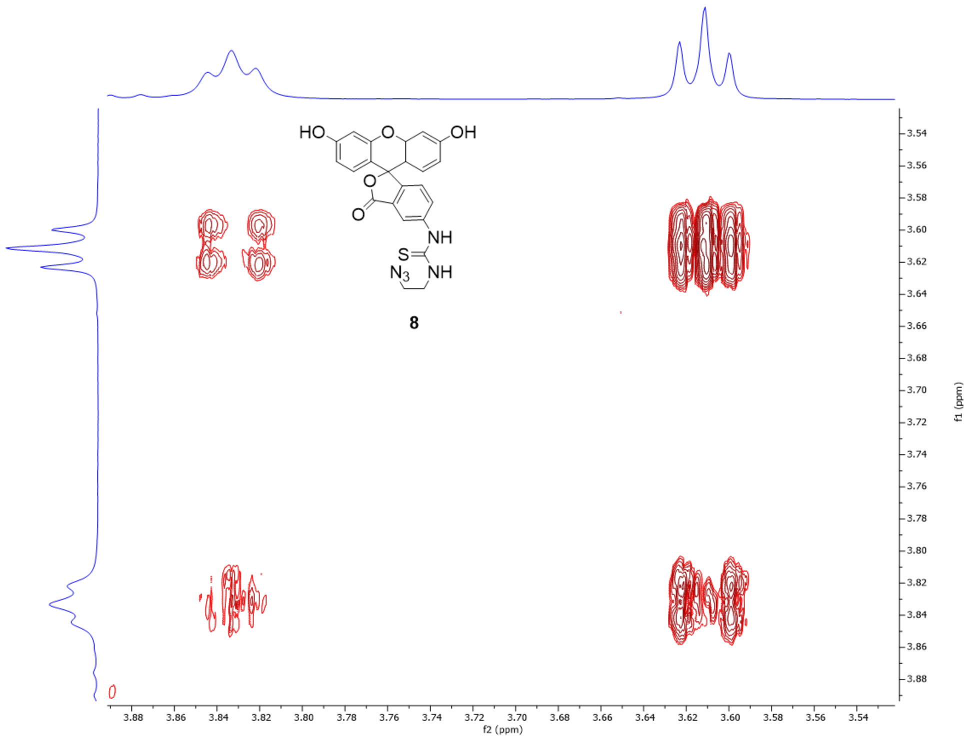

**Figure S34**. COSY spectrum of compound **8**.

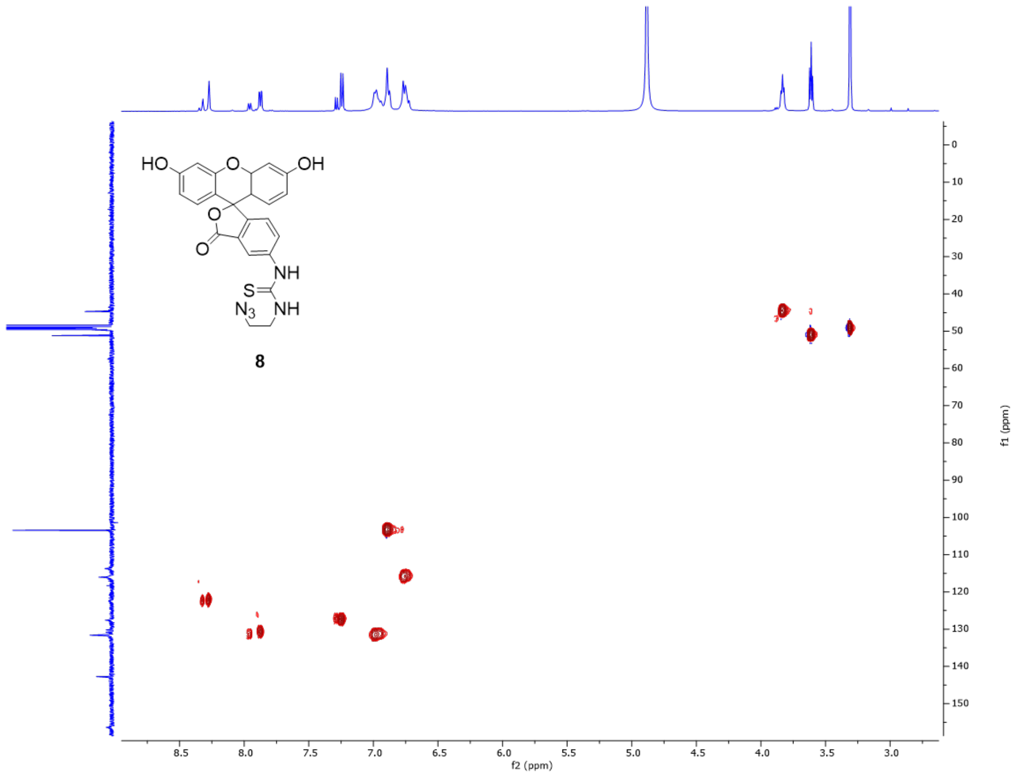

**Figure S35**. HSQC spectrum of compound **8**.

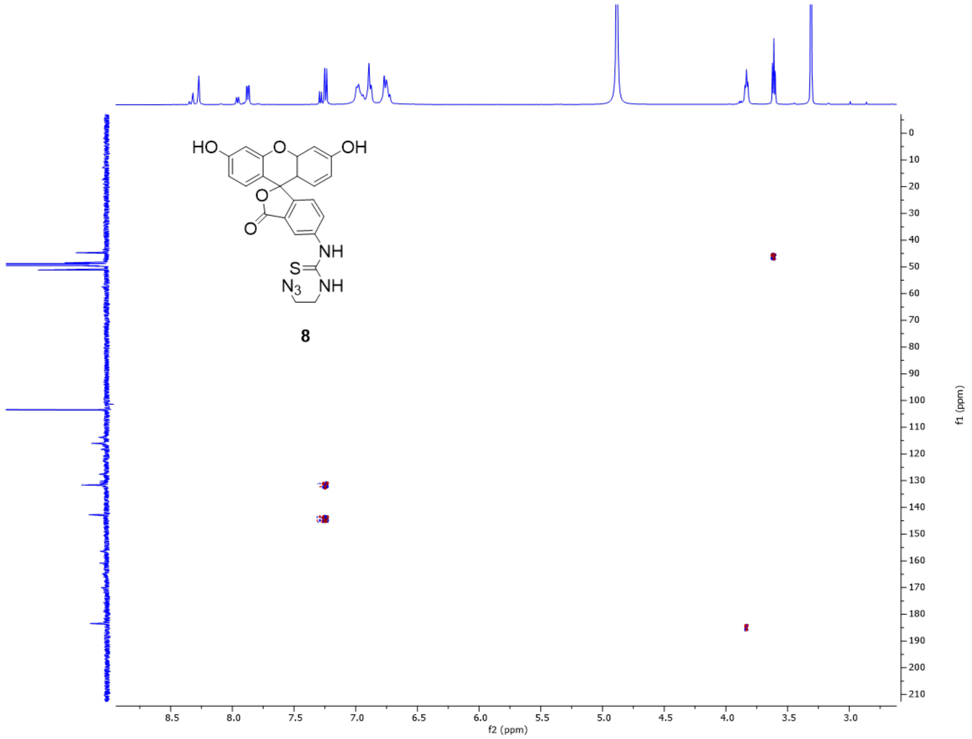

**Figure S36**. HMBC spectrum of compound **8**.

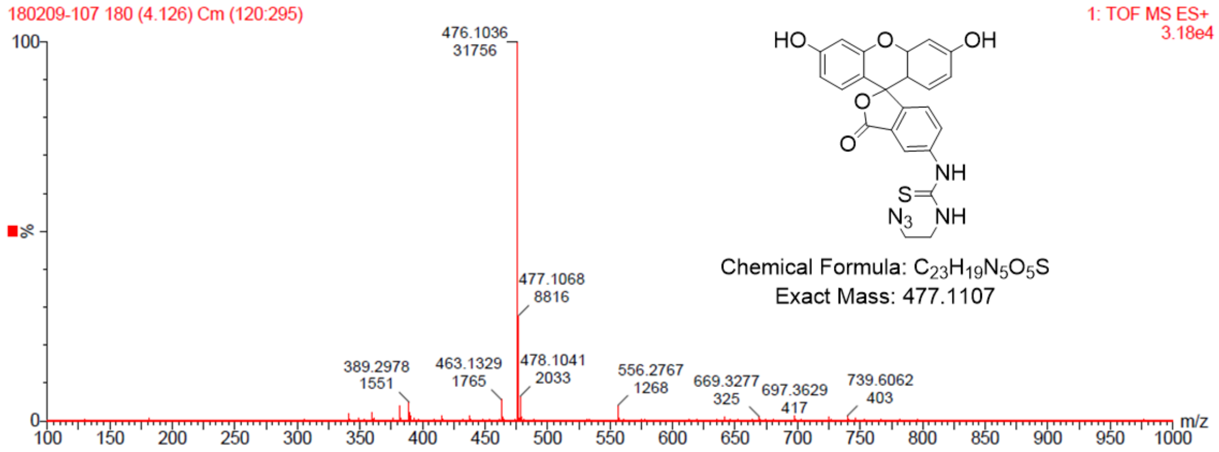

**Figure S37**. ESI-MS (positive mode) of compound **8**.

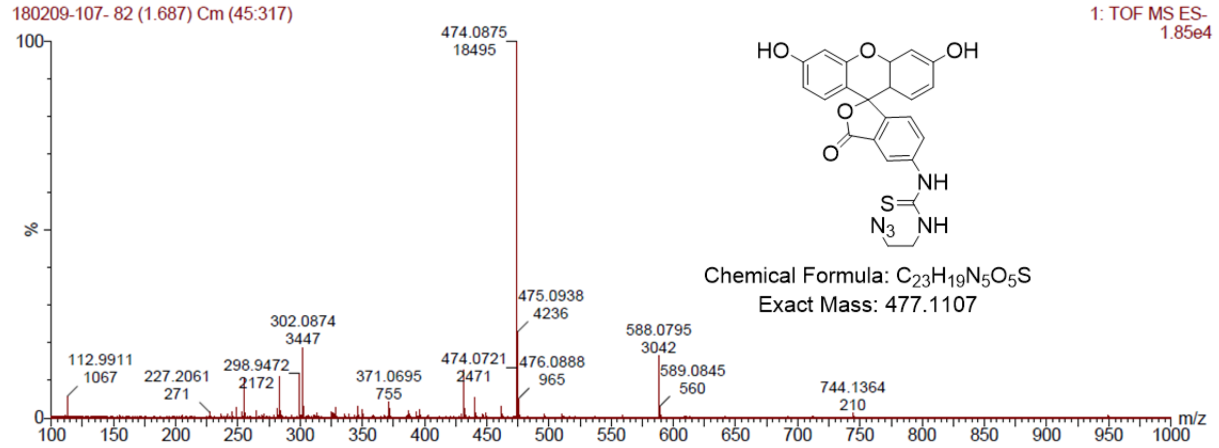

**Figure S38**. ESI-MS spectrum (negative mode) of compound **8**.

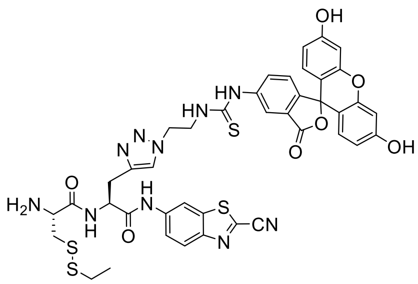

**1**

**(R)-2-amino-N-((S)-1-((2-cyanobenzo[d]thiazol-6-yl)amino)-3-(1-(2-(3-(3',6'-dihydroxy-3-oxo-3H-spiro[isobenzofuran-1,9'-xanthen]-5-yl)thioureido)ethyl)-1H-1,2,3-triazol-4-yl)-1-oxopropan-2-yl)-3-(ethyldisulfaneyl)propenamide (1):** Compounds **4** (5 mg, 0.0115 mmol, 1.0 eq.) and **6** (6 mg, 0.0115 mmol, 1.0 eq.) were dissolved in 600 uL DMF and added to an $N_2$-charged flask. $CuSO_4{\cdot}5H_2O$ (1.2 uL of 0.5 M aqueous solution, 0.05 eq.) added, then $(BimC_4A)_3$ (60 uL of 0.01 M aqueous sol'n, 0.05 eq.), then sodium ascorbate (60 uL of 1.0 M sol'n, 5.0 eq.). The resulting orange solution was covered in tin foil and stirred under $N_2$. After 2h, good conversion was observed by HPLC. Solution centrifuged and purified by HPLC to give the desired product as a yellow powder (2.2 mg, 22%). $^1$H-NMR (500 MHz, MeOD): δ/ppm = 8.59 (1 H, d, J 2.0), 8.10 (1 H, d, J 9.0), 8.01 (1 H, d, J 1.5), 7.88 (1 H, s), 7.68 – 7.61 (2 H, m), 7.16 (1 H, d, J 8.2), 6.70 (2 H, t, J 2.2), 6.66 (2 H, dd, J 8.4, 4.6), 6.53 (2 H, ddd, J 16.5, 8.7, 2.1), 4.98 – 4.92 (1 H, m), 4.68 (2 H, dt, J 13.8, 4.6), 4.29 (1 H, dd, J 8.6, 4.8), 4.17 – 4.00 (2 H, m), 3.43 – 3.38 (1 H, m), 3.35 – 3.33 (1 H, m), 3.29 – 3.24 (1 H, m), 3.04 – 2.96 (1 H, m), 2.77 – 2.61 (2 H, m), 1.25 (4 H, t, J 7.3). MS (ESI) calc'd for $C_{41}H_{36}N_{10}O_7S_4$: 907.1579 $[M-H]^-$, found 907.1584.

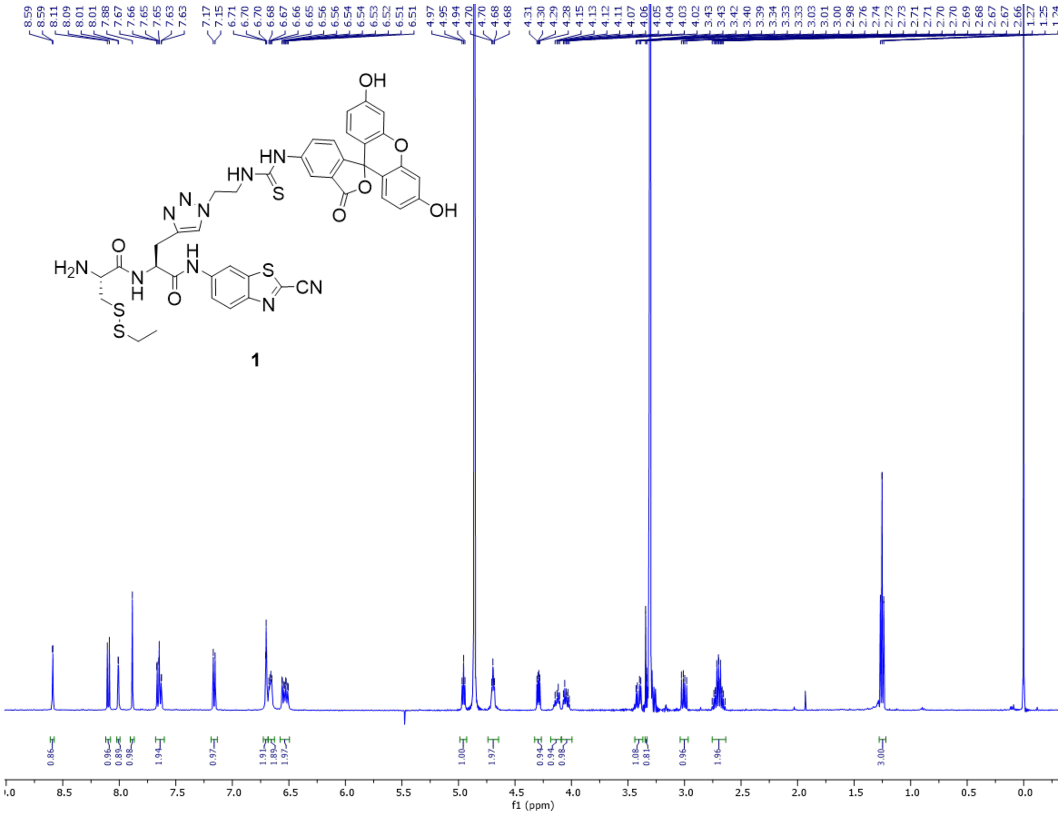

**Figure S39**. $^1$H-NMR spectrum of compound **1**.

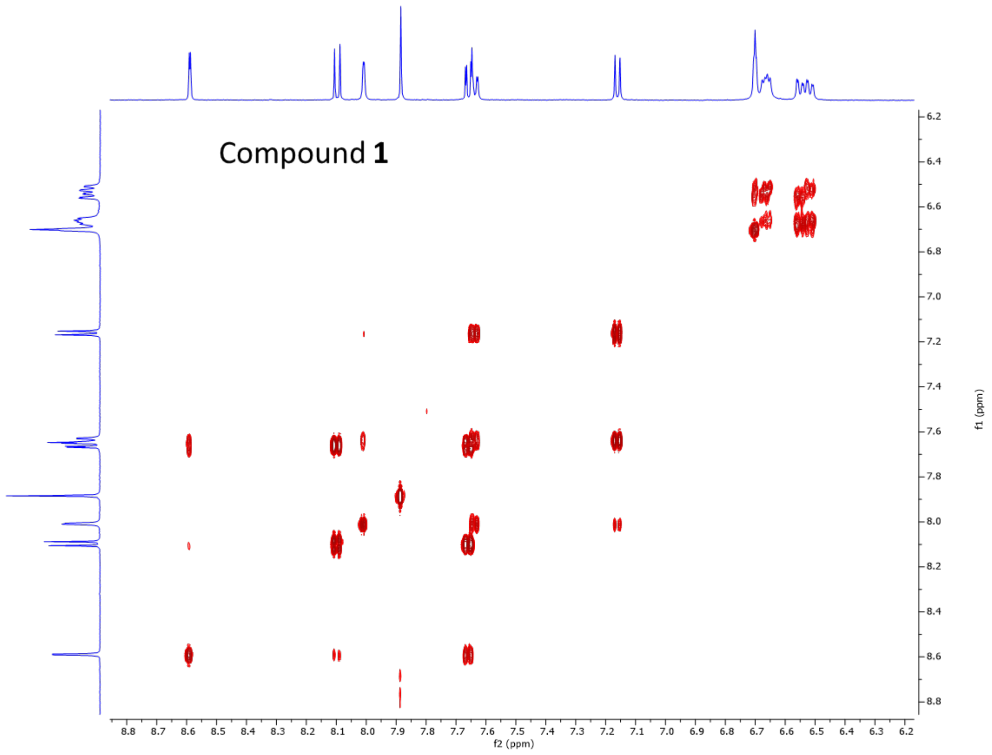

**Figure S40**. COSY spectrum of compound **1**.

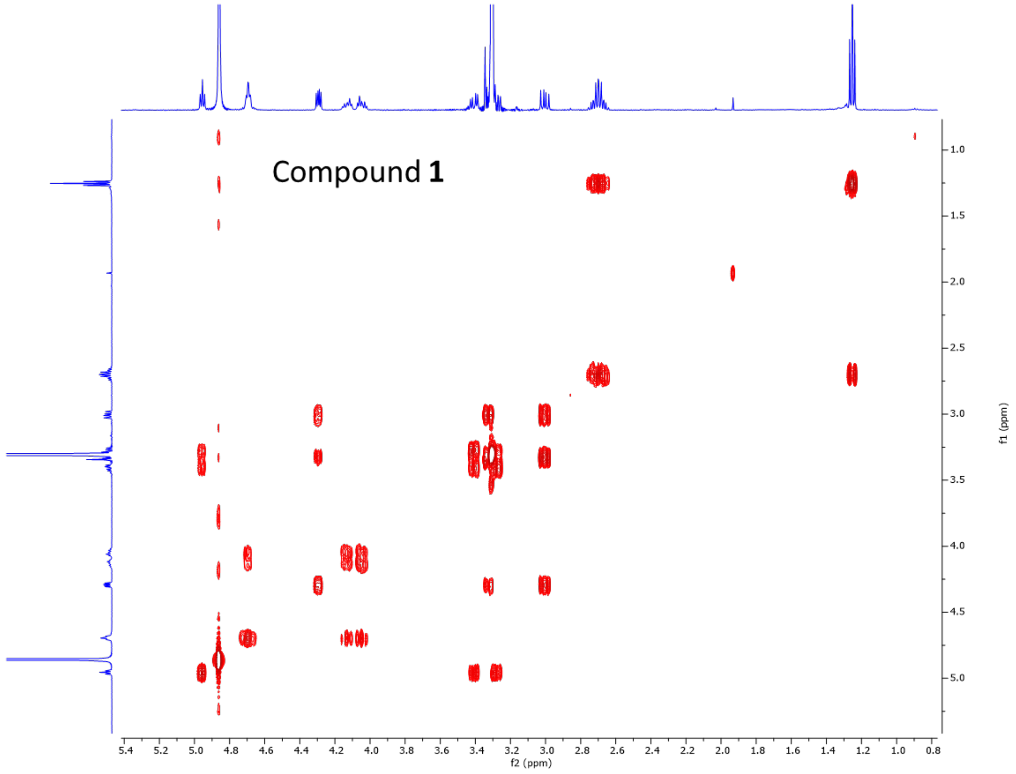

**Figure S41**. COSY spectrum of compound **1**.

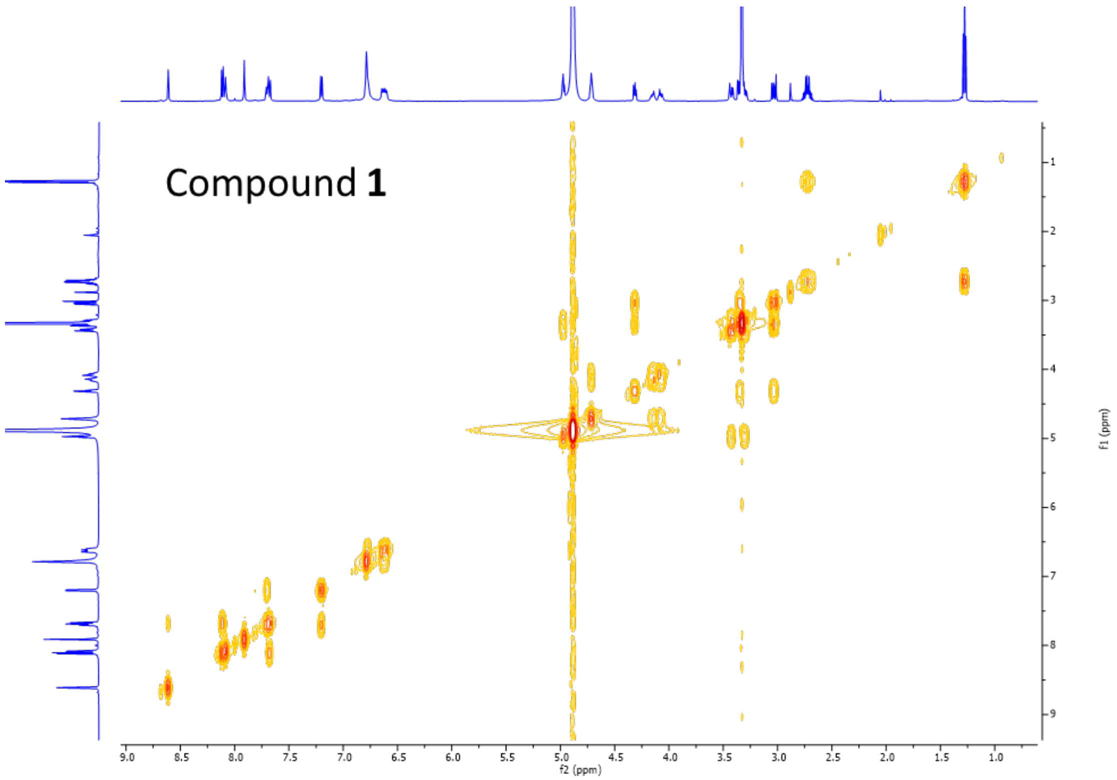

**Figure S42**. COSY spectrum (full) of compound **1**.

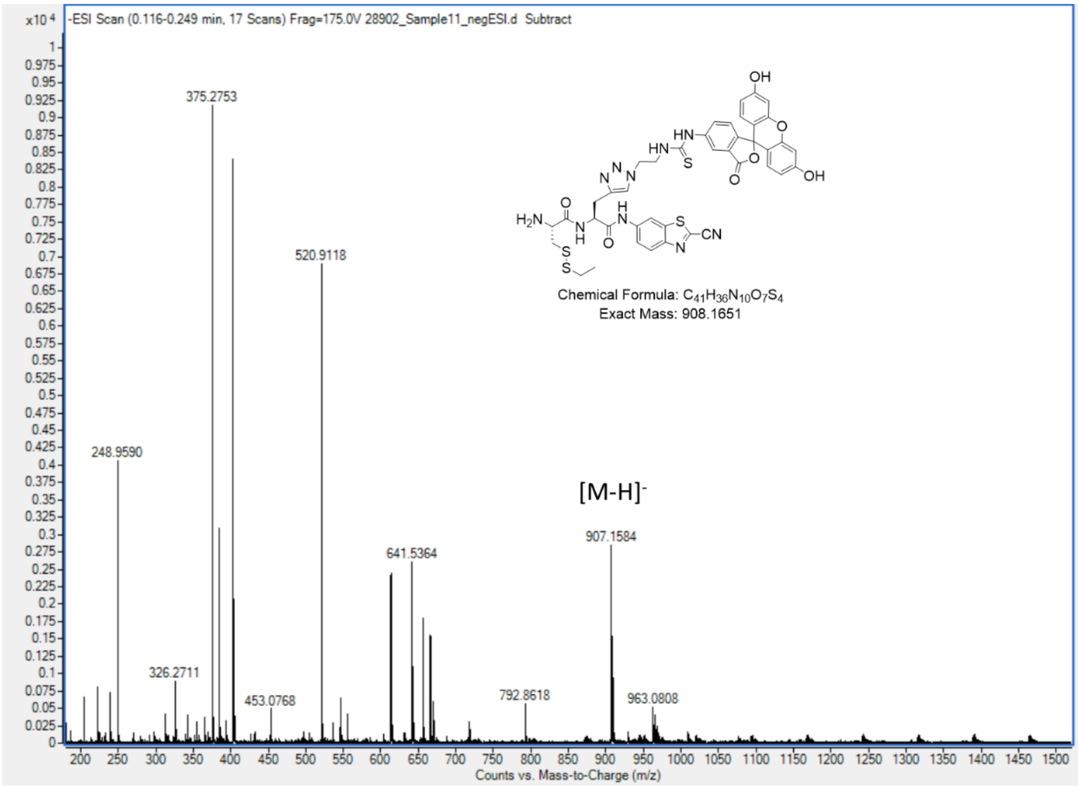

**Figure S43**. ESI-MS of compound **1**.

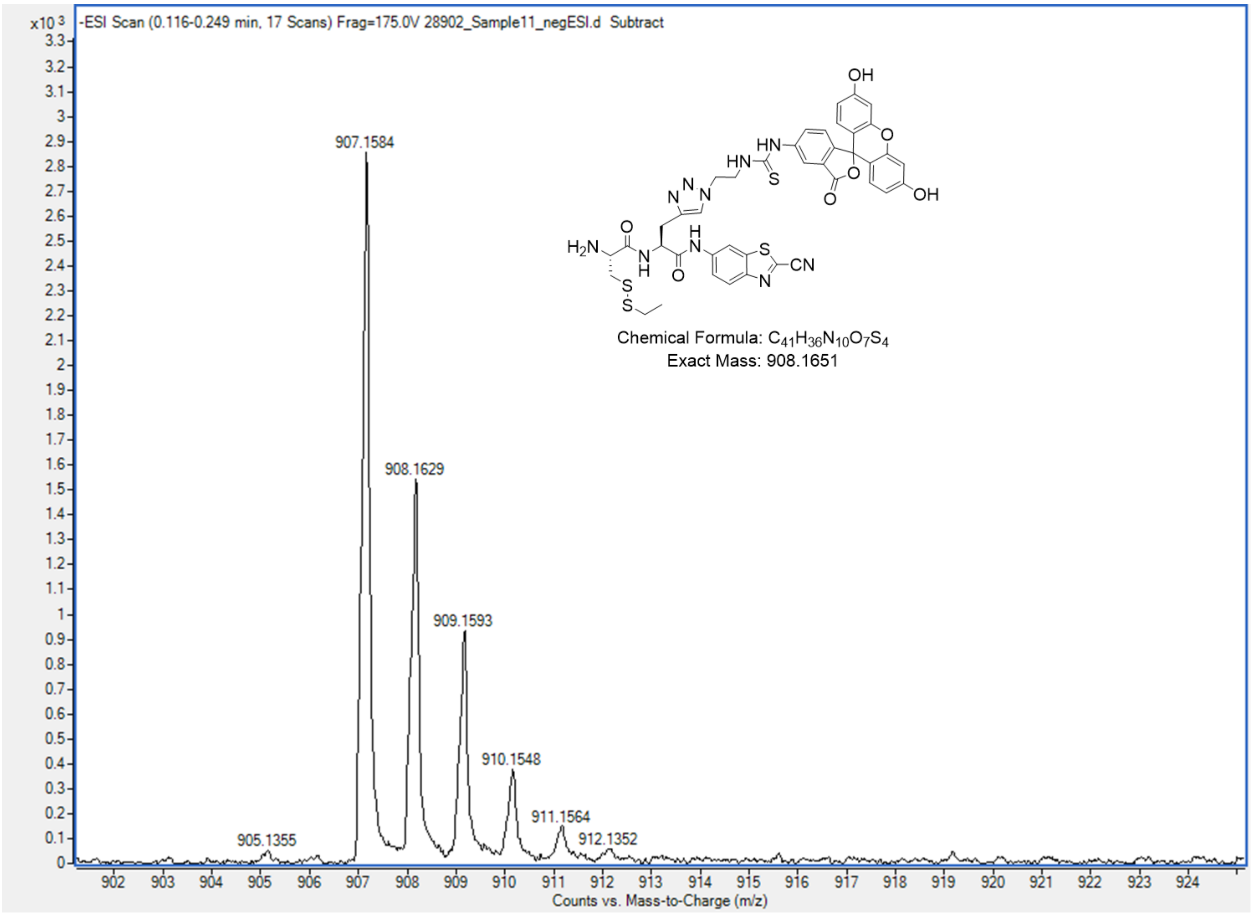

**Figure S44**. ESI-MS of compound **1**.

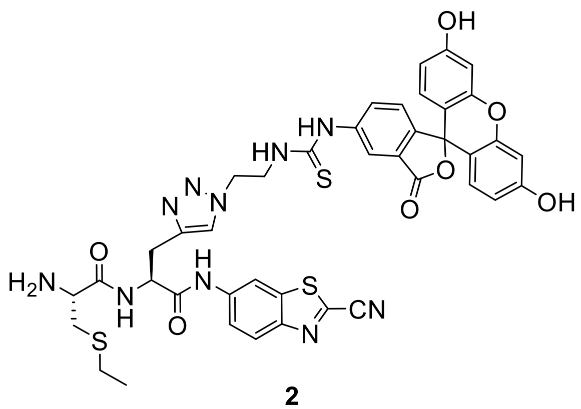

**2**

**(R)-2-amino-N-((S)-1-((2-cyanobenzo[d]thiazol-6-yl)amino)-3-(1-(2-(3-(3',6'-dihydroxy-3-oxo-3H-spiro[isobenzofuran-1,9'-xanthen]-5-yl)thioureido)ethyl)-1H-1,2,3-triazol-4-yl)-1-oxopropan-2-yl)-3-(ethylthio)propanamide (2):**

$^1$H NMR (600 MHz, Methanol-$d_4$) δ 8.51 (d, *J* = 2.1 Hz, 1H), 8.02 (d, *J* = 9.0 Hz, 1H), 7.90 (d, *J* = 2.0 Hz, 1H), 7.79 (s, 1H), 7.58 (dd, *J* = 9.0, 2.1 Hz, 1H), 7.54 (dd, *J* = 8.1, 2.0 Hz, 1H), 7.07 (d, *J* = 8.2 Hz, 1H), 6.59 (t, *J* = 2.4 Hz, 2H), 6.55 (dd, *J* = 8.7, 4.0 Hz, 2H), 6.43 (ddd, *J* = 15.5, 8.7, 2.4 Hz, 2H), 4.87 (t, *J* = 6.3 Hz, 3H), 4.63 – 4.58 (m, 2H), 4.03 (dq, *J* = 10.5, 5.3 Hz, 2H), 4.00 – 3.93 (m, 1H), 3.31 (d, *J* = 5.8 Hz, 1H), 3.06 (dd, *J* = 14.6, 5.0 Hz, 1H), 2.79 (dd, *J* = 14.6, 8.7 Hz, 1H), 2.50 (q, *J* = 7.4 Hz, 2H), 1.09 (t, *J* = 7.4 Hz, 3H), 0.81 (t, *J* = 6.9 Hz, 1H). MS (ESI) calc'd for $C_{41}H_{36}N_{10}O_7S_3$: 875.1858 [M-H]$^-$, found 875.1825.

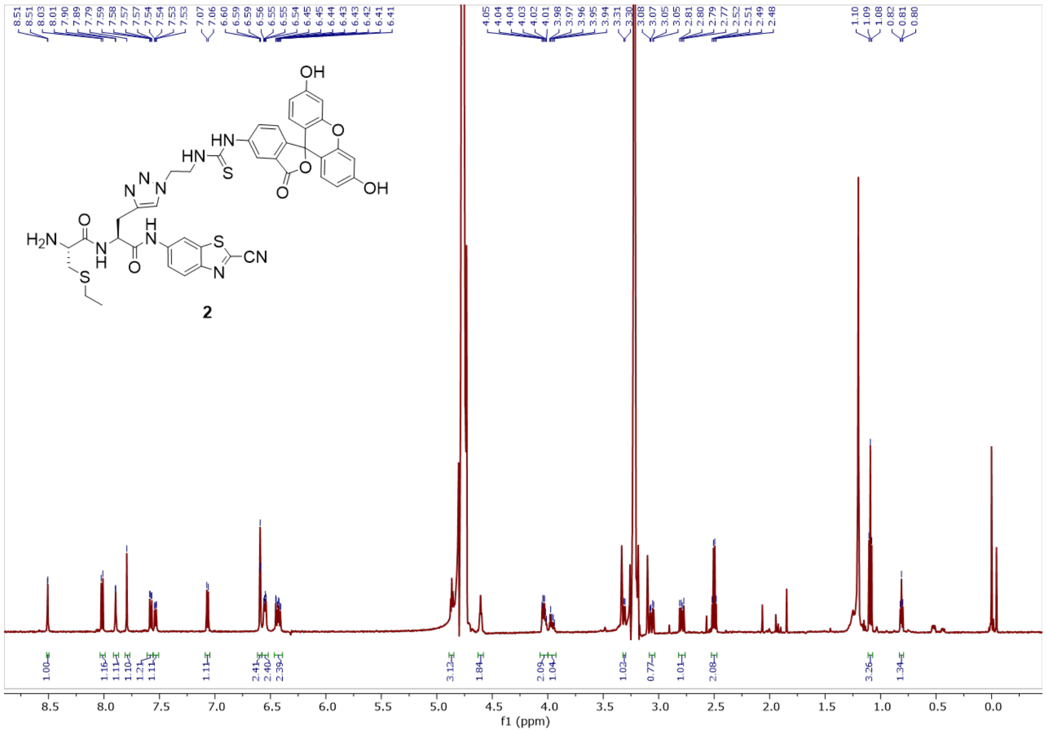

**Figure S45**. $^1$H-NMR spectrum of compound **2**.

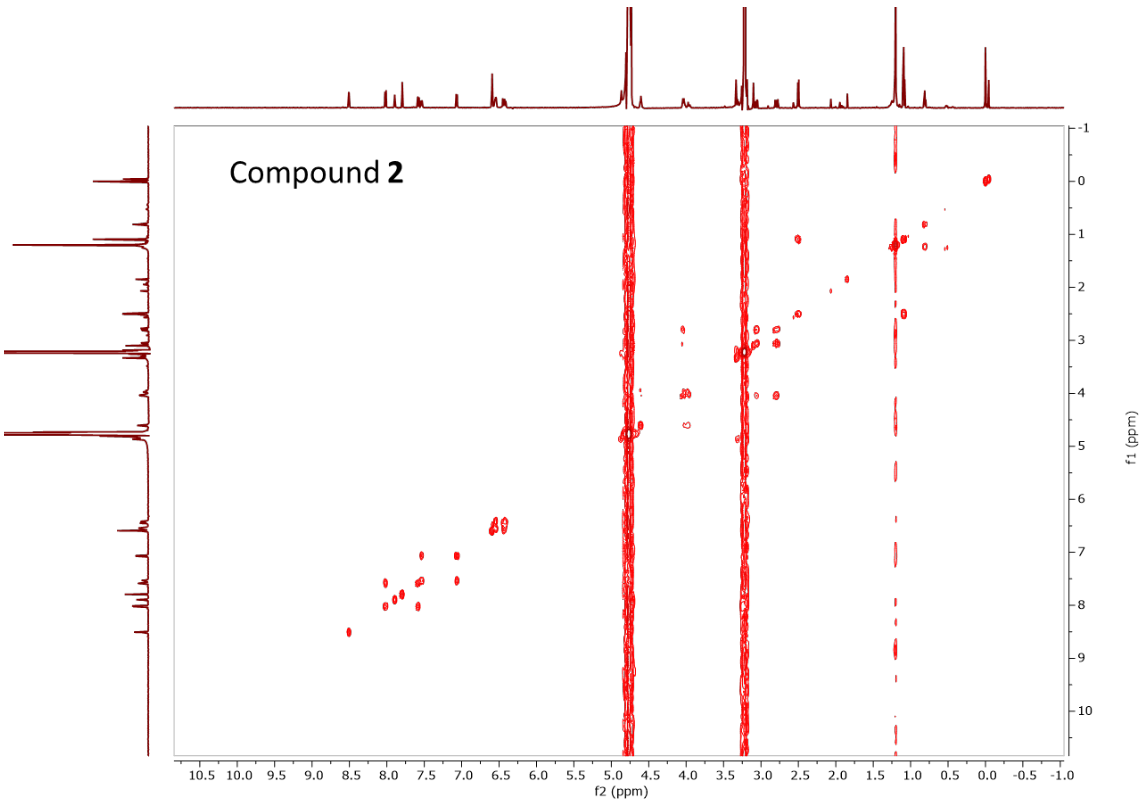

**Figure S46**. COSY spectrum of compound **2**.

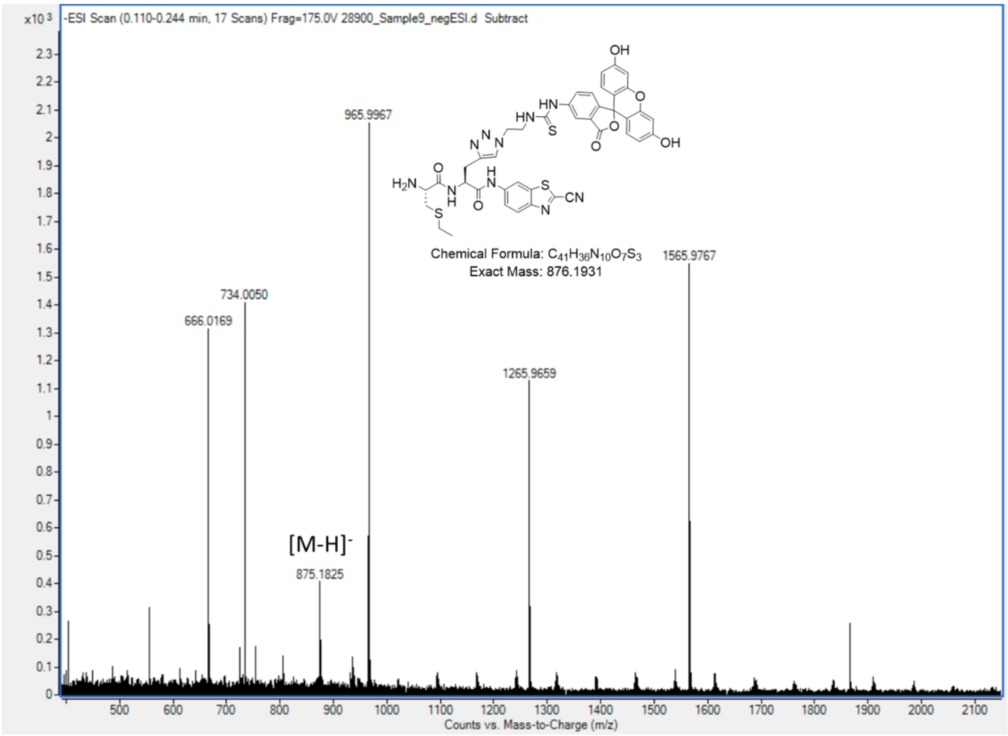

**Figure S47**. ESI-MS of compound **2**.

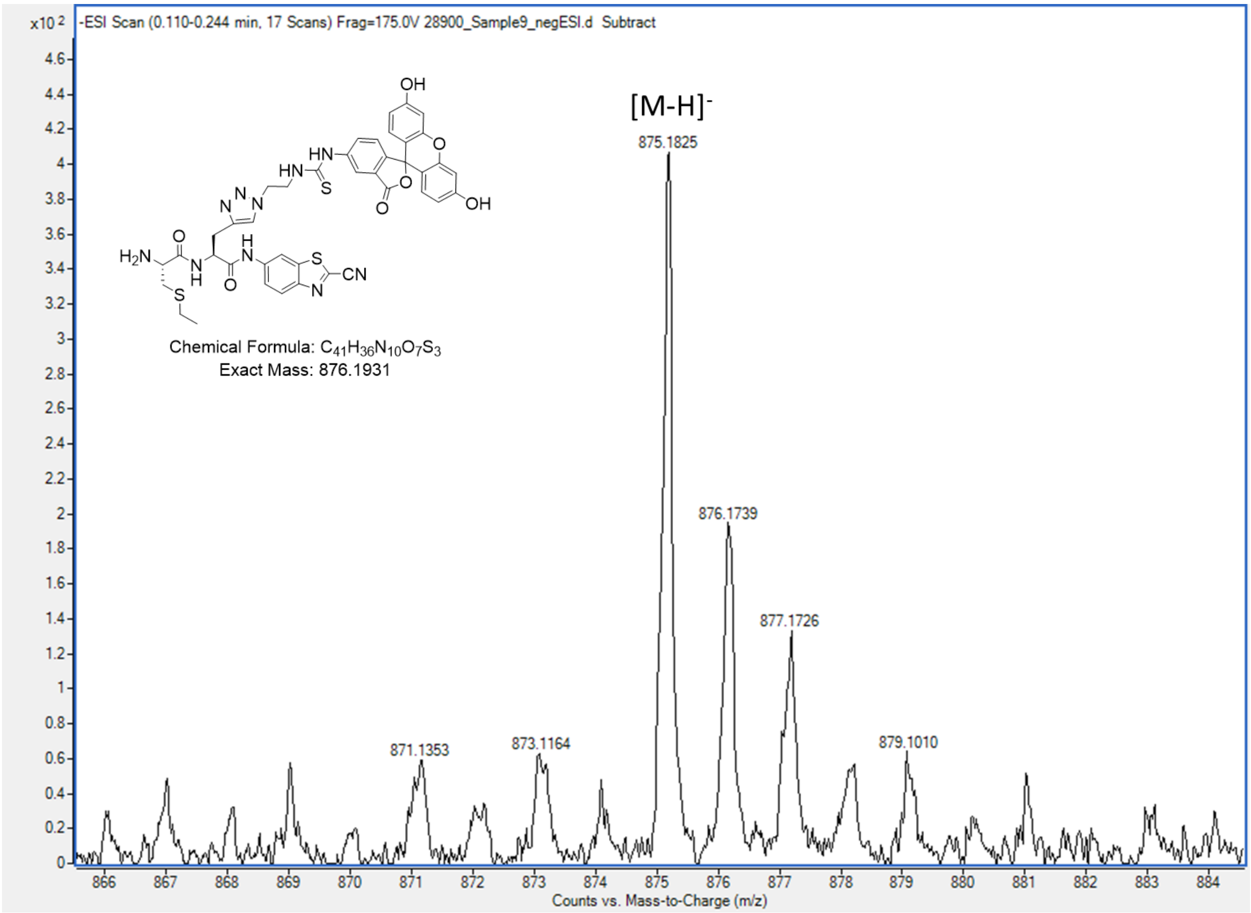

**Figure S48**. ESI-MS of compound **2**.
